# Hybridization during the adaptive radiation of *Oxera* (Lamiaceae) in New Caledonia: Is flower shape shift driven by introgression?

**DOI:** 10.1101/2025.06.07.658409

**Authors:** Ryusuke Ikeda, Ryoya Kawai, Gildas Gâteblé, Yoshihisa Suyama, Shun K. Hirota, Makoto Terauchi, Hideki Noguchi, Yuji Isagi

## Abstract

Recent genomic studies have suggested that hybridization may play a significant role in adaptive radiation, rapid speciation, and convergent evolution. The genus *Oxera*, a plant taxon thought to have diversified at its beginning through adaptive radiation in New Caledonia, provided an opportunity to investigate these processes. Within the *robusta* subclade of *Oxera*, characterized by bird-pollinated yellow-orange flowers, convergent evolution of flower shape is likely to have occurred. We aimed to elucidate the hybridization history of the *robusta* subclade by whole genome sequencing and MIG-seq data. Our analyses revealed an ancestral introgression from *O. coriacea* to *O. sympatrica*, whose flowers are remarkably similar to each other. Among the introgressed genomic regions, we identified several genes potentially involved in flower shape development. *O. sympatrica* and its sympatric sister species exhibit distinct flower shapes, and pollinator-mediated reproductive isolation presumed to be a major barrier between them. The ancestral introgression uncovered in this study may have driven the convergent evolution of flower shape in the *robusta* subclade and played a crucial role in the speciation process of *O. sympatrica*. These finding contribute to our understanding of the interplay between hybridization, adaptive radiation, and speciation process.

## Introduction

Hybridization is an important evolutionary process that can facilitate both adaptation and speciation. The evolutionary consequences of hybridization are diverse, including population fusion, reinforcement of reproductive isolation, and formation of allopolyploids (Runemark et al., 2019). One notable consequence is introgression - the transfer of genetic variation from one population to another through hybridization followed by repeated backcrosses (Anderson & Hubricht, 1938). When the introgressed genetic variation provides an adaptive advantage in the recipient population, it may increase in frequency and fix in the recipient genome. Such adaptive introgression can lead to convergent evolution between donor and recipient lineages (Stern, 2013). In some taxa, it has been shown that introgression had driven the repeated evolution of an ecologically important trait, such as the wing colour patterns of *Heliconius* butterflies (The Heliconius Genome Consortium, 2012) and the flower colour of *Mimulus* monkey flowers (Stankowski & Streisfeld, 2015). As seen in adaptive introgression, hybridization can expand the range of gene pools and combinations of genetic variation without relying on *de novo* mutation. Hence, hybridization has been hypothesized to contribute to adaptive radiation and rapid speciation (Marques et al., 2019; Seehausen, 2004). This creative role of hybridization may be particularly important in adaptive radiation on islands, where genetic diversity is typically constrained by bottleneck and inbreeding following colonization (Cerca et al., 2023).

New Caledonia is recognized as one of the major biodiversity hotspots where explosive diversification of several taxa has taken place. Although New Caledonia is a continental island that separated from Gondwana, its history of submergence (Palaeocene to Eocene) and re-emergence (Oleocene), combined with subsequent isolation until today, has endowed it with characteristics similar to those of oceanic islands, contributing to biotic disharmony and high levels of regional endemism (Grandcolas et al., 2008). Regarding vascular plants, approximately 75% of the more than 3,400 species inhibiting on the island are endemic. Certain plant taxa in New Caledonia are considered examples of adaptive radiation due to their notable morphological and ecological diversity alongside significant species richness (Cerca et al., 2023). Investigating the role of hybridization in these adaptive ra diations could provide insights into the evolutionary processes that have shaped New Caledonia’s diverse flora. However, studies examining hybridization in New Caledonian plant taxa remain limited and are often based on analysis of only a few genetic loci (Pillon et al., 2009; Keppel et al., 2011). An exception is Paun et al. (2016), which utilized RAD-seq data to demonstrate that interspecies hybridization occurred during the adaptive radiation of the genus *Diospyros* (Ebenaceae). This study suggests that hybridization may have played a limited role in the repeated soil habitat shifts during the adaptive radiation of *Diospyros*. However, whether similar patterns apply to other ecological traits or to other adaptive radiations of New Caledonian plant taxa remains uncertain. Furthermore, comprehensive studies using whole-genome data have yet to be conducted.

The genus *Oxera* (Lamiaceae) is a plant taxon that has predominantly diversified in New Caledonia. Unlike many other plant taxa in the island, this genus exhibits high diversification not only in habitat preferences but also in life form and fruit and flower morphologies, making it an example of island adaptive radiation according to Cerca et al. (2023). In New Caledonia, the genus comprises more than 30 species grouped into 8 phylogenetic subclades (Barrabé et al., 2019). The *robusta* subclade, in particular, primarily includes robust liana species with bird-pollinated large yellow-orange flowers. Within this subclade, two distinct flower types are recognized (fig.1): The first type is characterized by a corolla tube that expands into a funnel shape at the apex, with elongated stamens and pistils that prominently protrude. The second type exhibits a corolla tube of almost uniform diameter from base to apex, with short stamens and pistils that only slightly extend beyond the tube. Hereafter, in this paper, the former will be referred to as the “funnel-type” and the latter as the “pipe-type”. Field observations indicate that these flower types may differ in the positioning of pollen placement on and effectiveness in pollination by two bird species, *Glycifohia undulata* and *Philemon diemenensis* (Gildas *et al*., unpublished data).

**Fig. 1.**
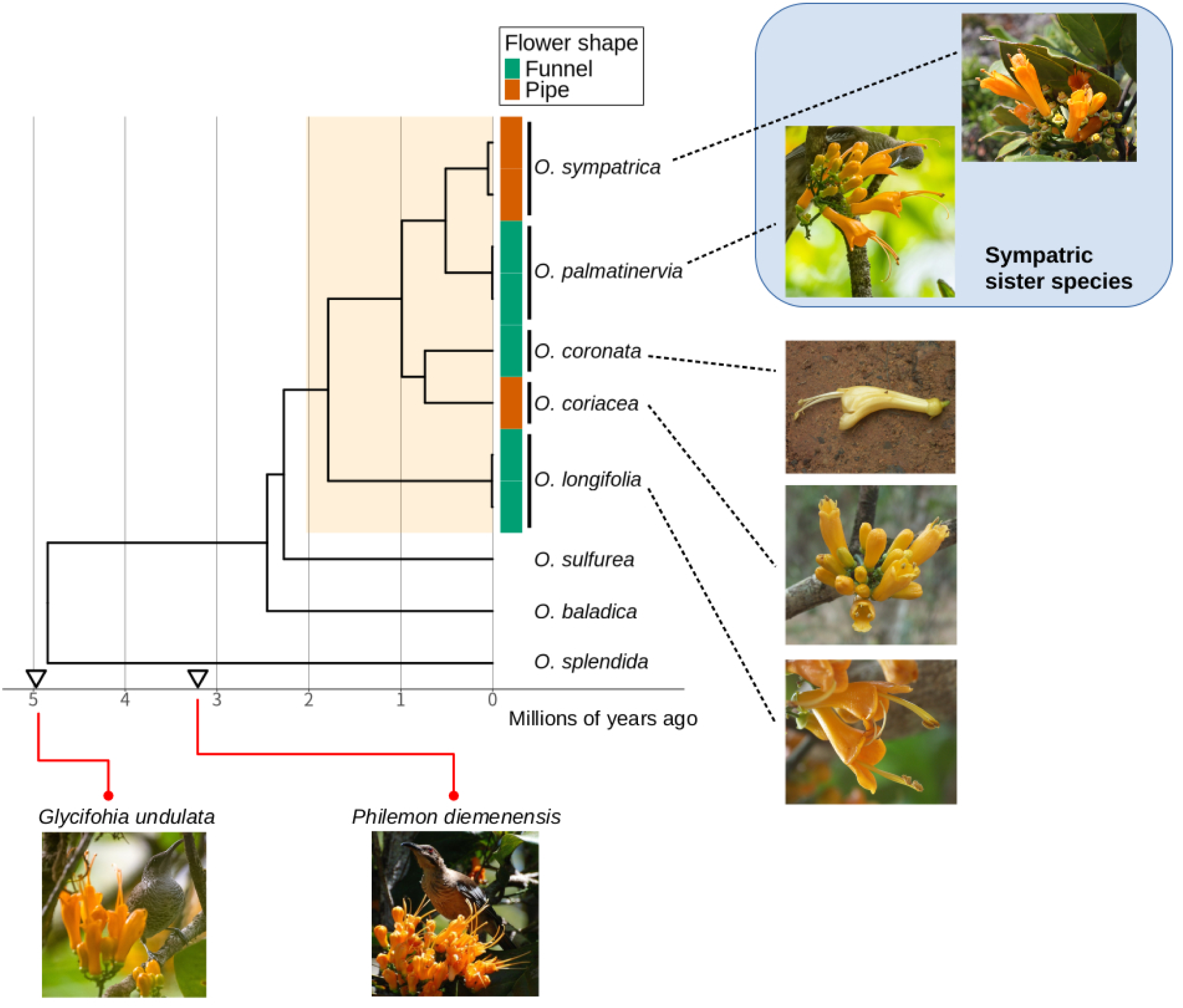
Time calibrated phylogenetic tree of the *robusta* subclade. Time calibrated tree estimated by SNAPP. The *robusta* subclade is highlighted in orange. The colors next to tips correspond to flower shape. The funnel-type is green and the pipe-type is brown. The inverted triangles on x-axis represent the stem ages of New Caledonia endemic pollinator species, *Glycifohia undulata* and *Philemon diemenensis*, estimated in Marki et al. (2017). The data were derived from TimeTree 5 (Kumar et al., 2022).

A recently described species in the *robusta* subclade, *O. sympatrica*, and its putative sister species, *O. palmatinervia*, coexist sympatrically in rainforests on ultramafic soils in the southern part of the main island, Grande Terre. While these two species exhibit similar vegetative and fruit morphologies, they differ in their flower type: *O. sympatrica* has the pipe-type flower, whereas *O. palmatinervia* has the funnel-type flower (fig.1). The recognition of *O. sympatrica* suggests that convergent evolution of flower shape may have occurred within the *robusta* subclade. In particular, the pipe-type flower species, *O. sympatrica* and *O. coriacea*, are not monophyletic, and each has a putative sister species with funnel-type flowers. To date, there has been no study on the hybridization history within *Oxera*, and the potential role of hybridization in the repeated flower shape evolution in the *robusta* subclade has not been investigated.

In this study, we assembled draft genomes of *O. balansae* from *balansae* subclade and *O. baladica* from *baladica* subclades, and employed whole-genome sequencing and the MIG-seq method (Suyama & Matsuki, 2015) to assess the phylogenetic relationships and hybridization history of the *robusta* subclade. Phylogenetic analyses based on whole-genome SNP data confirmed that *O. sympatrica* is a sister species to *O. palmatinervia* and revealed that multiple shifts in flower shape have occurred within the subclade. Our population structure analysis and hybridization inferences consistently suggested that introgression has occurred from *O. coriacea* to *O. sympatrica*, both of which possess pipe-type flowers. Additionally, we examined genomic local allele sharing and phylogenetic relationships to identify introgressed genomic regions shared between *O. sympatrica* individuals. Our findings indicate that the similar flower shape observed between non-sister species in the *robusta* subclade may have evolved through ancestral introgression. These results contribute to a deeper understanding of the role of hybridization in island adaptive radiations and its influence on the diversification of New Caledonia’s rich flora.

## Materials and Methods

### Sampling and Genome assembly

All plant materials sequenced in this study were collected from fields and Insitut Agronomique Néo-Calédonien Research Station of Saint-Louis (IAC) between 2006 and 2020, with the exception of *O. splendida*. For *O. splendida*, the DNA sample extracted in Barrabé et al. (2015), Crayn1217, was used.

We selected one sample of *O. baladica* (YI-1351) and one sample of *O. balansae* (YI-1350) for draft genome assembly. For *O. balansae*, the DNA sample was extracted using the CTAB method (Doyle & Doyle, 1987) and sequenced on GridION (Oxford Nanopore, Oxford, UK) and Hiseq X (Illumina, San Diego, CA, USA) at 150bp paired-end. Adapter trimming and quality filter of Illumina short-read were conducted by Trimmomatic 0.38 (Bolger et al., 2014) with ILLUMINACLIP:2:30:10, AVGQUAL:10, and MINLEN:20. Assembly was performed by NECAT 0.0.1 (Y. Chen et al. 2021) with Nanopore long-read. The obtained genome assembly was polished using Racon 1.3.1 (Vaser et al. 2017) with long-read. After five repeated runs of Racon, the assembly was further polished using Pilon 1.23 (Walker et al. 2014) with Illumina short-read. Then, redundant contigs due to the heterozygosity were identified and removed using Purge Haplotigs (Roach et al. 2018). The total length of the assembly was 514,803,507 bp and the number of contigs was 1,085. The N50 was 1,381,998 bp.

For *O. baladica*, the DNA sample was extracted with NucleoBond HMW DNA (Macherey-Nagel, Düren, Germany) and sequenced on Sequel II (Pacific Biosciences, California, USA) and Hiseq 2000 (Illumina, San Diego, CA, USA) at 250 bp paired-end. Adapter trimming and quality filter of Illumina short-read were conducted by FASTP (S. Chen, 2023). First, we performed two assemblies by different assemblers, FALCON/FALCON-UNZIP (Chin et al., 2016) and FLYE (Kolmogorov et al., 2019), with PacBio long-read. Those two assemblies were then processed by PURGE_DUPS (Guan et al., 2020) to remove duplicate contigs and then merged by quickmerge (Chakraborty et al., 2016). The merged assembly was polished two times by NextPolish (Hu et al., 2020) with Illumina short-read. The repeat regions were masked using repeatmodeler2 (Flynn et al., 2020) and repeatmasker (Smit et al., 2013-2015). Since some protein sequences could be incorrectly detected as repeat in these processes, putative protein-coding repeats were identified and unmasked using custom Perl program for generating the masked genome sequence and the repeat annotation file in RepeatMasker output format. The total length of the assembly was 491,814,862 bp and the number of contigs and scaffolds was 356. The N50 was 6,742,774 bp.

We annotated the draft genome of *O. baladica* using RNA-seq data of matured leaf and flower from 11 samples of 8 species (2 samples from *O. doubetiae*, *O. neriifolia*, and *O. pancheri*; 1 sample from *O. baladica*, *O. garoensis*, *O. longifolia*, *O. microcalyx*, and *O. pulchella*). Total RNA was extracted with Agilent Plant RNA Isolation Mini Kit (Agilent Technologies, Santa Clara, CA, USA) from fixed tissues by RNAlater (Thermo Fisher Scientific, San Jose, CA, USA). RNA samples derived from single individuals were sequenced on Illumina Hiseq 2000 platform at 100 bp paired-end. FASTP was used for adapter trimming and quality filter. Transcripts were assembled by Trinity (Grabherr et al., 2011) and mapped to the draft genome of *O. baladica* to get the information of exon, CDS, and UTR regions by PASA pipeline (Haas et al., 2003). Transcripts with shared exons were grouped into one gene using custom shell script. Then, final genome annotation was conducted with GINGER (Taniguchi et al., 2023). In the homology-based module of GINGER, the protein database was built from eight land plants including Lamiaceae species, *Arabidopsis thaliana* (https://ftp.ncbi.nlm.nih.gov/genomes/all/GCF/000/001/735/GCF_000001735.4_TAIR10.1/GCF_000001735.4_TAIR10.1_protein.faa.gz), *Solanum lycopersicum* (http://ftp.ensemblgenomes.org/pub/release-53/plants/fasta/solanum_lycopersicum/pep/Solanum_lycopersicum.SL3.0.pep.all.fa.gz), *Oryza sativa* (http://ftp.ensemblgenomes.org/pub/release-53/plants/fasta/oryza_sativa/pep/Oryza_sativa.IRGSP-1.0.pep.all.fa.gz), *Sesamum indicum* (https://ftp.ncbi.nlm.nih.gov/genomes/all/GCF/000/512/975/GCF_000512975.1_S_indicum_v1.0/GCF_000512975.1_S_indicum_v1.0_protein.faa.gz), *Perilla frutescens* PF040 (https://figshare.com/ndownloader/files/38016120), *Perilla frutescens* PC002 (https://ftp.ncbi.nlm.nih.gov/genomes/all/GCA/019/512/045/GCA_019512045.2_ICMM_Pcit_2.0/GCA_019512045.2_ICMM_Pcit_2.0_protein.faa.gz), *Salvia splendens* (https://ftp.ncbi.nlm.nih.gov/genomes/all/GCF/004/379/255/GCF_004379255.1_SspV2/GCF_004379255.1_SspV2_protein.faa.gz), *Salvia hispanica* (https://ftp.ncbi.nlm.nih.gov/genomes/all/GCF/023/119/035/GCF_023119035.1_UniMelb_Shisp_WGS_1.0/GCF_023119035.1_UniMelb_Shisp_WGS_1.0_protein.faa.gz). The alignment of the protein sequences against the genome by SPALN was performed with the “Angiosp” parameter file. In the RNA-seq-based module of GINGER (mapping-base/*de novo*-base), one sample of *O. baladica* described above was used. In the merge step of GINGER, the gene models from homology-, RNA-seq-and *ab initio*-based modules were merged with that of PASA described above as the external evidence. Using the obtained proteins as queries, DIAMOND BLASTP searches (Buchfink et al., 2015) were performed against the protein database (uniref90), and domain information were obtained from INTERPROSCAN (Jones et al., 2014). As a result, we obtained 28333 coding-genes. The evaluation by BUSCO (Manni et al., 2021) against the database of embryophyta (embryophyta_odb10, n=1614, proteins mode) showed the good completeness (C:95.6%[S:87.0%, D:8.6%], F:0.6%, M:3.8%).

### Sequencing and genotyping of whole-genome resequencing data set

A total of 13 samples from 8 species were resequenced (supplementary table S1). Genomic DNA was extracted using either the CTAB method (Doyle & Doyle, 1987) or DNeasy Plant Mini kit (Qiagen, Hilden, Germany) and 150 bp paired-end reads were sequenced on the Hiseq X or Novaseq 6000 platform (Illumina, San Diego, CA, USA). Adapter trimming and read filtering were executed using Trimmomatic 0.39 (Bolger et al., 2014) with ILLUMINACLIP:2:30:10, AVGQUAL:10, and MINLEN:20. Paired reads were aligned to the *O. baladica* draft genome using BWA-mem 0.7.17-r1188 (Li, 2013) with-K 100,000,000, and the output was sorted by SAMtools 1.15.1 (Danecek et al., 2021) sort command. Duplicate reads were removed by MarkDuplicates command in GATK 4.1.9 (McKenna et al., 2010) with--OPTICAL_DUPLICATES_PIXCEL_DISTANCE 2,500 and--REMOVE_DUPLICATES true. Considering only reads with mapping quality >= 30 and bases with quality >= 20, reference cover rates were 0.72-0.99 and means of depth were 10.9-50.1. For *O. splendida* (splend1), the non-New Caledonian outgroup, reference cover rate was 0.72 and mean depth was 50.1.

Variant calling was performed by freebayes-parallel script in FreeBayes 1.3.6 (Tange, 2011; Garrison & Marth, 2012) with--use-mapping-quality,--genotype-qualities,-m 30,-q 20, and--populations. Mean depth per site per individual was calculated with VCFtools 0.1.16 (Danecek et al., 2011), and sites with a mean depth per individual greater than twice the average depth were excluded using BCFtools 1.13 (Danecek et al., 2021) view command with-e SUM(FORMAT/DP)/(N_SAMPLES-N_MISSING) > 48. Then, sites with QUAL score less than 30 and genotypes with depth under 5 were filetered out using GATK’s VariantFiltration and SelectVariants commands with –exclude-filtered and-set-filtered-gt-to-nocall. Called haplotypes were decomposed into SNPs and indels using the vcfdecompose command in RTG Tools 3.12.1 (Cleary et al., 2015) with--break-mnps and only SNPs were extracted with the view command in BCFtools with-V indels,mnps,other. Using vcflib 1.0.3 (Garrison et al., 2022) vcffixup command, alternate allele count (AC), alternate allele frequency (AF), and number of called (NS) in the INFO field were re-calculated, then invariant sites across all individuals were filtered out by BCFtools view command with-i ‘AC>0’ and-e ‘AF=1’. After all, 63,348,015 variant sites were used for downstream data handling.

### Phylogenetic analysis based on whole-genome scale SNPs

We used the SNP concatenation and multispecies coalescent approaches to infer the phylogenetic relationship within the *robusta* subclade. For the SNP concatenation approach, we used IQ-TREE 1.6.12 (Nguyen et al., 2015) to reconstruct the ML tree based on non-missing and missing < 20% SNP datasets. Site filtering and format converting were performed by vcf2phylip.py 2.8 (Ortiz, 2019). For each dataset, ML inference was performed with the GTR+G+ASC model, and 1,000 replicates for ultrafast bootstrap (Hoang et al., 2018) and SH-like approximate likelihood ratio test (SH-aLRT) (Guindon et al., 2010) were performed. The non-missing dataset consisted of 18,448,714 sites and the missing < 20% dataset consisted of 27,217,566 sites. For the multispecies coalescent approach, we used SVDQ (Chifman & Kubatko, 2014) implemented in PAUP* 4.0a169 (Swofford, 2003). SVDQ was performed based on 1kb, 5 kb, and 10 kb apart SNP datasets without missing data. Non-missing SNP sites were kept by BCFtools (Danecek et al., 2021) view command and thinned to 1 kb, 5 kb, and 10 kb apart by VCFtools (Danecek et al., 2011)--thin. Each dataset was converted to nexus format by vcf2phylip.py (1 kb-dataset 322,723 sites, 5 kb-dataset 76,171 sites, 10 kb-datasets 40,208 sites). For each run, 500 bootstrap replicates were performed.

### Chloroplast genome assembly and bayesian divergence time estimation

To reveal the timeframe of the *robusta* subclade evolution, we performed bayesian divergence time estimations based on chloroplast genome sequences and genome-wide SNP. First, we assembled the chloroplast genomes of *O. baladica* (baladi1) and *O. splendida* (splend1) and used BEAST 2.6.7 (Bouckaert et al., 2019) to estimate the crown age of *Oxera* based on the fossil records in Lamiaceae. Then, the divergence time within the *robusta* subclade was estimated based on the BEAST2 result and genome-wide SNP using SNAPP 1.6.1 (Bryant et al., 2012; Stange et al., 2018).

Chloroplast genome sequences were assembled by GetOrganelle 1.7.6.1 pipeline (Camacho et al., 2009; Bankevich et al., 2012; Langmead & Salzberg, 2012; Jin et al., 2020) and/or NOVOPlasty 4.2.1 (Dierckxsens et al., 2016) and polished by Pilon 1.24 (Walker et al., 2014). To include some calibration nodes in our analysis, complete chloroplast genome sequences of 11 species from 9 genera in Lamiaceae were downloaded from NCBI based on phylogenetic trees of previous studies (supplementary table S2) (Roy & Lindqvist, 2015; Zhao et al., 2021; Rose et al., 2022). The chloroplast genome sequences were split into LSC, SSC and IR, aligned by MAFFT 7.508 (Katoh & Standley, 2013) with FFT-NS-i algorithm, and trimmed by trimAl 1.4 (Capella-Gutiérrez et al., 2009). ModelFinder (Kalyaanamoorthy et al., 2017) implemented in IQ-TREE 1.6.12 (Nguyen et al., 2015) was used to perform BIC-based substitution model selection for each region. The crown ages of the subfamily Nepetoideae and the *Stachys* s.l. clade (Stachydeae excluding *Melittis*) were calibrated based on the fossil records, and the crown age of Lamiaceae were calibrated based on the results of Rose et al. (2022). Using BEAST 2.6.7, we conducted three independent runs of Markov chain Monte Carlo (MCMC) with a chain length of 300,000,000, sampling every 10,000 generations. MCMC convergence was checked using Tracer 1.7.2 (Rambaut et al., 2018), and the log files and tree files were combined using LogCombiner 2.6.7 with a burn-in of 10%. The Maximum Clade Credibility tree was generated by TreeAnnotator 2.6.2. In addition, a ML phylogenetic estimation was performed using IQ-TREE to confirm the divergences around the calibration points. More details on the assembly and phylogenetic inference of chloroplast genomes are written in the Supplementary Methods.

Using SNAPP 1.6.1 implemented in BEAST 2.7.1, we estimated the divergence times among the *robusta* subclade species based on genome-wide SNP and the BEAST2 result. As SNAPP is computationally demanding, the samples used for the analysis were restricted to 11 individuals from 8 species ( *O. splendida:* splend1, *O. baladica*: baladi1, *O. sulfurea*: sulfur1, *O. longifolia*: longif1 and robust1, *O. coronata*: corona1, *O. coriacea*: coriac1, *O. palmatinervia*: palmat2-3, and *O. sympatrica*: sympat1-2). Biallelic sites were extracted with genomics_general 0.4 (https://github.com/simonhmartin/genomics_general). SNP sites, 70 kb apart from each other, were extracted by VCFtools (5,853 SNPs). The vcf was converted to phylip by vcf2phylip 2.9 (Ortiz, 2019) and the XML configuration file was written by snapp_prep.rb (Stange et al., 2018). Based on the estimated crown age of *Oxera* by BEAST2 with chloroplast genome sequences (mean 5.88 MYA, median 5.33 MYA, 95% confidence interval 1.66–11.52 MYA), the crown age of *Oxera* was constrained by a log-normal distribution (mean in real space 5.88, standard deviation in log space 0.45). We conducted two independent runs of MCMC with a chain length of 10,000,000, sampling every 5000 generations. MCMC convergence was checked using Tracer 1.7.2, and the log files and tree files were combined using LogCombiner 2.7.7 with a burn-in of 10%. The Maximum Clade Credibility tree was generated by TreeAnnotator 2.7.7 and visualized by ggtree 3.10.0 (Yu et al., 2017).

### Hybridization history inference

Multispecies coalescent networks were estimated using SNaQ (Solís-Lemus & Ané, 2016) implemented in Phylonetworks 0.15.3 (Solís-Lemus et al., 2017). Non-missing SNP sites were retained using the BCFtools 1.13 (Danecek et al., 2021) view command, and SNPs separated by 2 kb were extracted by VCFtools 0.1.16 (Danecek et al., 2011). The VCF file was subsequently converted to PHYLIP format using the vcf2phylip.py 2.8 (Ortiz, 2019). Concordance factors were calculated for all possible 715 quartets of the 13 individuals using SNPs2CF 1.5 (Olave & Meyer, 2020). The average number of SNPs used for calculation of the concordance factor per quartet was 9182.1. The network estimation was performed using SnaQ. At hmax = 0, the 5kb-apart SVDQ tree was used for the starting topology. Then, the estimated topology at hmax = 0 was used for the starting topology for subsequent inferences (hmax >= 1). For each inference, 100 independent network searches were performed.

Dsuite 0.5 r49 (Malinsky et al., 2021) was used to calculate Patterson’s D (Patterson et al., 2012), admixture fraction f (Durand et al., 2011; Martin et al., 2015), and f-branch statistic (Malinsky et al., 2018) based on the whole genome SNPs. The 5 kb-apart SVDQ tree was used as an input tree for the calculation of those statistics with *O. splendida* as the outgroup. Dsuite DtriosParallel command was used to calculate D-statistic and f fraction for all possible trios of individuals. Genome sequences were partitioned into blocks by 40,000 informative SNPs and the jackknife method was applied using the-j 40000 option. To estimate gene flow in ancestral lineages, f-branch statistics were computed from the inferred f fraction using Dsuite Fbranch command. Only trios with p-values below the Bonferroni correction threshold of 0.05 were considered for the calculation of f-branch statistics.

### Population structure analysis based on Mig-seq

Our analyses showed that ancient introgression from *O. coriacea* to *O. sympatrica* had occurred (See Results section). Genome-wide SNP analysis was conducted using multiplexed inter-simple sequence repeat genotyping by sequencing (MIG-seq) method (Suyama & Matsuki, 2015). For MIG-seq analysis, we used 65 *O. palmatinervia*, 24 *O. sympatrica*, and 22 *O. coriacea* (supplementary table S3). The MIG-seq library was prepared following the two-step PCR method described by Suyama et al. (2022) and sequenced using an MiSeq system (Illumina, San Diego, CA, USA) with 80 bp paired-end reads. The initial 17 bases (comprising SSR and anchor regions) in both forward and reverse reads were excluded using the “DarkCycle” option in MiSeq Control Software (Illumina).

Primer sequences and low-quality reads were removed using Trimmomatic 0.39 (Bolger et al., 2014) with the parameters ILLUMINACLIP:2:30:10, SLIDINGWINDOW:4:15, and MINLEN:80. This filtering process yielded 20,840,104 reads (an average of 187,749 ± 4588 reads per sample) from an initial 22,813,860 raw reads (an average of 205,530 ± 4960 reads per sample). The draft genome of *O. balansae* was used as a reference for SNP mapping and identification. Quality-controled reads were aligned to the reference genome using BWA 0.7.17 (Li, 2013) with default parameters. Alignments were converted from SAM format to sorted and indexed BAM files using SAMtools 1.7 (Danecek et al., 2021). Genotypes were called using gstacks program of Stacks 2.53 (Catchen et al., 2013; Rochette et al., 2019) with default settings. Only SNPs present in at least 80% of samples were retained, and the sites showing excess heterozygosity (> 0.6) were excluded. For genetic structure analyses, the minimum allele frequency threshold was set to 5%, and only the first SNP per locus was included, resulting in 1,013 retained SNPs.

The population genetic structure was analyzed using STRUCTURE (Pritchard et al., 2000) with 20 independent runs, each with a burn-in of 100,000 steps followed by 100,000 MCMC steps under the admixture model. The log-likelihoods for each number of cluster (*K* = 1–10) were estimated, and optimal *K* values was determined using the Delta *K* method of (Evanno et al., 2005) in Structure Harvester (Earl & VonHoldt, 2012). Graphical results were obtained using CLUMPAK (Cluster Markov Packager Across K) (Kopelman et al., 2015).

### Genomic regions affected by introgression

To explore the introgressed genomic regions from *O. coriacea* that are shared between *O. sympatrica* individuals from Yaté and Dzumac (sympat1 and sympat2), sliding window analysis of fd statistic (Martin et al., 2015) and Saguaro (Zamani et al., 2013) were performed for whole-genome resequencing data. The fd statistic estimates the introgressed region between P2 and P3 under a combination (((P1, P2), P3), O). Using ABBABABAwindows.py from the genomics_general 0.4 (https://github.com/simonhmartin/genomics_general), the fd and D statistics were calculated in non-overlap 10 kb windows for the following four-taxon combinations to estimate the introgressed regions between *O. coriacea* and *O. sympatrica* individuals. *O. splendida* as an outgroup and 3 *O. palmatinervia* individuals as one population, (1) P1 = *O. palmatinervia*, P2 = *O. sympatrica* from Yaté (sympat1), P3 = *O. coriacea* (coriac1), (2) P1 = *O. palmatinervia*, P2 = *O. sympatrica* from Dzumac (sympat2), P3 = *O. coriacea* (coriac1), (3) P1 = *O. coronata* (corona1), P2 = *O. coriacea* (coriac1), P3 = *O. sympatrica* from Yaté (sympat1), (4) P1 = *O. coronata* (corona1), P2 = *O. coriacea* (coriac1), P3 = *O. sympatrica* from Dzumac (sympat2). Windows with fewer than 100 informative SNPs were considered as missing (-m 100). Based on the minimum estimated f-branch value between *O. sympatrica* and *O. coriacea* (see the Results section), windows with fd statistic in the top 2.6 % (1282 windows) were selected for each combination. As the fd statistic does not offer meaningful values when the BABA pattern is excessive (gene flow between P1 and P3) (Martin et al., 2015), only windows with a positive D statistic were considered. Windows commonly extracted in (1)–(4) were considered as candidate introgressed windows.

Saguaro is a program to segment aligned genome sequences into blocks with different phylogenetic relationships, by generating “cactus”, a matrix representing differences between sequences, and assigning it to genomic regions. 9 individuals from 6 taxa were used for the analysis (*O. sulfurea:* sulfur1, *O. longifolia:* longif1, *O. coronata:* corona1, *O. coriacea:* coriac1, *O. palmatinervia:* palmat1-3, and *O. sympatrica:* sympat1-2). After the targeted samples were extracted, sites with more than four called individuals were extracted using BCFtools 1.13 (Danecek et al., 2021). Conversion from the vcf file to the feature file was performed by VCF2HMMFeature and sites with no variation among samples were removed. The process of cactus assignment and generation of a new cactus was repeated 15 times and the updating and re-assignment of existing cactus was repeated 5 times in each process. Each cactus was converted to phylip format by Saguaro2Phylip and a neighbor-joining tree was estimated by rapidNJ (Simonsen et al., 2008). As cactus13 is consistent with the expected introgression genealogy (fig. 5), the fragments to which cactus13 was assigned were considered as candidate regions.

Finally, we explored genes affected by the introgression from *O. coriacea* to *O. sympatrica* ancestral lineage. Cactus13 segments overlapped with candidate introgressed 10kb-windows of fd statistic were extracted by BEDtools 2.31.1 (Quinlan & Hall, 2010) and genes overlapped these segments were listed.

## Results

### Phylogenetic placement of O. sympatrica

The SNP concatenation ML trees and the multispecies coalescent trees were almost identical and most branches had high support values. The ML trees estimated by IQ-TREE based on two datasets, non-missing SNP sites and missing < 20% SNP sites, had full support values of SH-LaRT and UFB for all branches and the same topology (supplementary fig. S2). The multispecies coalescent trees estimated by SVDQ based on three datasets, 1 kb-apart SNPs, 5 kb-apart SNPs, and 10 kb-apart SNPs, had high support values (> 90) and the same topology, except for the intra-species relationships within *O. longifolia* (supplementary fig. S3). In all phylogenetic trees, the inter-species topology was completely the same, in agreement with previous studies (Barrabé et al., 2019). *O. sympatrica*, a recently described new species, is inferred to be sister to *O. palmatinervia* with full support values in all analyses. Two pipe-type flower species, *O. sympatrica* and *O. coriacea*, were estimated to be non-monophyletic, and their sister species have funnel-type flowers.

### Divergence time estimation based on chloroplast genome and genome-wide SNPs

The ML and Bayesian phylogenetic inferences based on the chloroplast genome sequence dataset of Lamiaceae resulted in consistent phylogenetic trees (supplementary fig. S4 and S5). Each branch had full support in SH-LaRT, UFB, and posterior probability (PP). BEAST2 estimated the average crown age of *Oxera* to be about 5.9 Mya (median: 5.3 Mya, 95% HPD: 1.7–11.5 Mya). SNAPP based on genome-wide SNPs resulted in a reliable phylogenetic tree with high PP (= 0.99 or more) for all branches ( fig. 1 and supplementary fig.S6). The SNAPP tree was highly consistent with the ML trees by IQ-TREE and multispecies coalescent trees by SVDQ. In SNAPP, the crown age of *Oxera* was estimated to be about 4.9 Mya (median: 4.4 Mya, 95% HPD: 1.4-9.8 Mya), and the crown age of the *robusta* subclade was estimated to be about 1.8 Mya (median: 1.6, 95% HPD: 0.5-3.6). These results are almost consistent with Barrabé et al. (2019), although the crown age of *Oxera* in SNAPP is slightly more recent. *O. sympatrica* and *O. palmatinervia* were estimated to be the most recently diverged species pair in this study (average: 0.5, median:0.5, 95% HPD: 0.1-1.0).

### Inference of hybridization history in the robusta subclade

Phylogenetic networks with hybrid nodes were estimated by SnaQ based on 2kb-apart non-missing SNPs. The phylogenetic networks were estimated up to hmax = 3, as there were only 2 reticulation nodes at hmax = 3. Pseudo-likelihood scores showed the largest improvement between hmax = 0 and hmax = 1, with only limited score improvements after hmax = 1 (fig.2 and supplementary fig.S8). At hmax=1, it was estimated that *O. sympatrica* is a hybrid lineage between *O. palmatinervia* and *O. coriacea*. The estimated inheritance probability *γ* from *O. coriacea* was 0.116, and 0.884 from *O. palmatinervia*.

**Fig. 2.**
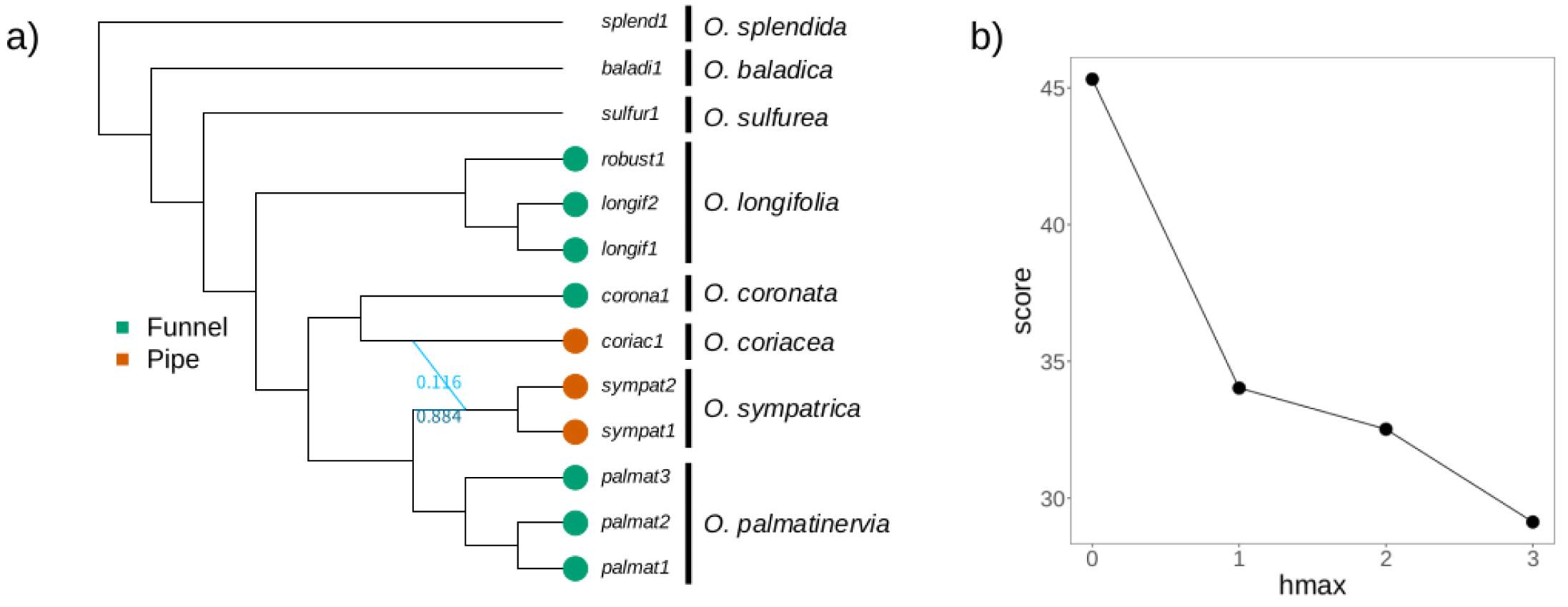
The SnaQ network (hmax=1) based on 2kb apart SNPs. (a) Blue lines represent hybrid edges. The numbers near the hybrid edges are inheritance probabilities which represent the proportion of genes inherited from each parental population. (b) The line plot show pseudo-likelihood score of each network (hmax=0-3).

D and f-branch statistics detected hybridization signals consistent with the phylogenetic networks by SNaQ. Significant D statistics were found between *O. sympatrica* and *O. coriacea* samples in all possible trios (Table.1). In f-branch statistic, high admixture proportions were detected between *O. sympatrica* and *O. coriacea* (fig. 3). Between *O. coriacea* (coriac1) and *O. sympatrica* from Yate and Dzumac (sympat1 and sympat2), f-branch were 0.043 and 0.042 (f_b=coriac1_(P3=sympat1 and sympat2)). Between *O. coriacea* and the ancestral lineage of *O. sympatrica* samples, f-branch was 0.026 (f_b=ancestral_ (P3=coriac1)).

**Table. 1.**
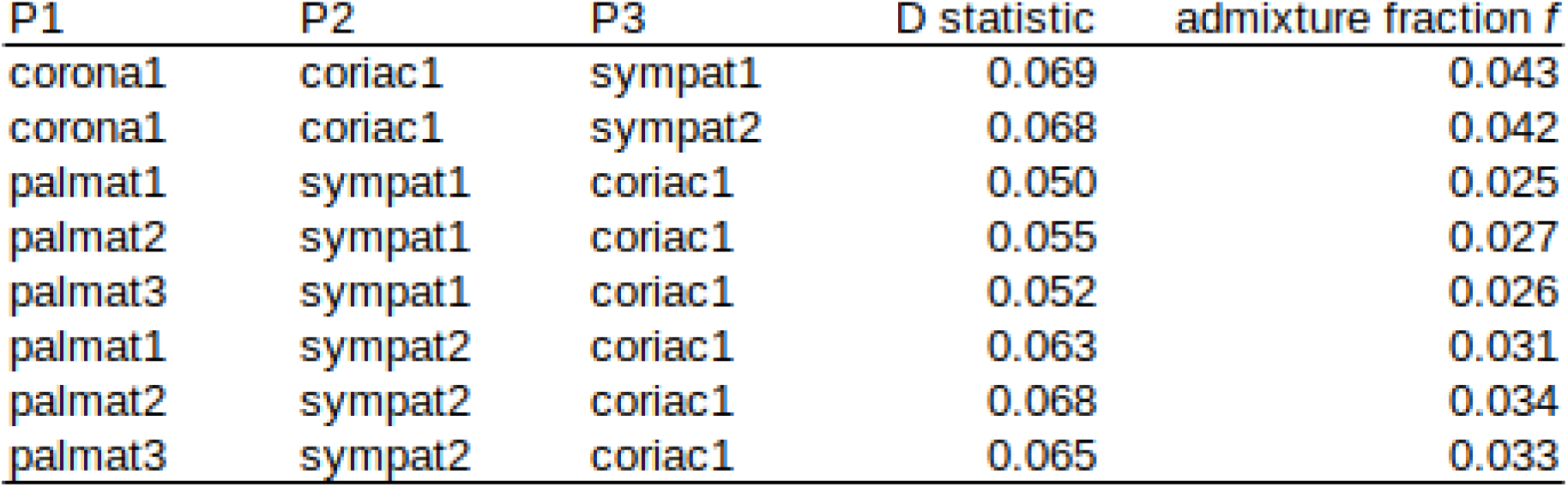
D statistic and admixture fraction f between *O. sympatrica* and *O. coriacea*.

| P1 | P2 | P3 | D statistic | admixture fraction $f$ |
| --- | --- | --- | --- | --- |
| corona1 | coriac1 | sympat1 | 0.069 | 0.043 |
| corona1 | coriac1 | sympat2 | 0.068 | 0.042 |
| palmat1 | sympat1 | coriac1 | 0.050 | 0.025 |
| palmat2 | sympat1 | coriac1 | 0.055 | 0.027 |
| palmat3 | sympat1 | coriac1 | 0.052 | 0.026 |
| palmat1 | sympat2 | coriac1 | 0.063 | 0.031 |
| palmat2 | sympat2 | coriac1 | 0.068 | 0.034 |
| palmat3 | sympat2 | coriac1 | 0.065 | 0.033 |

**Fig. 3.**
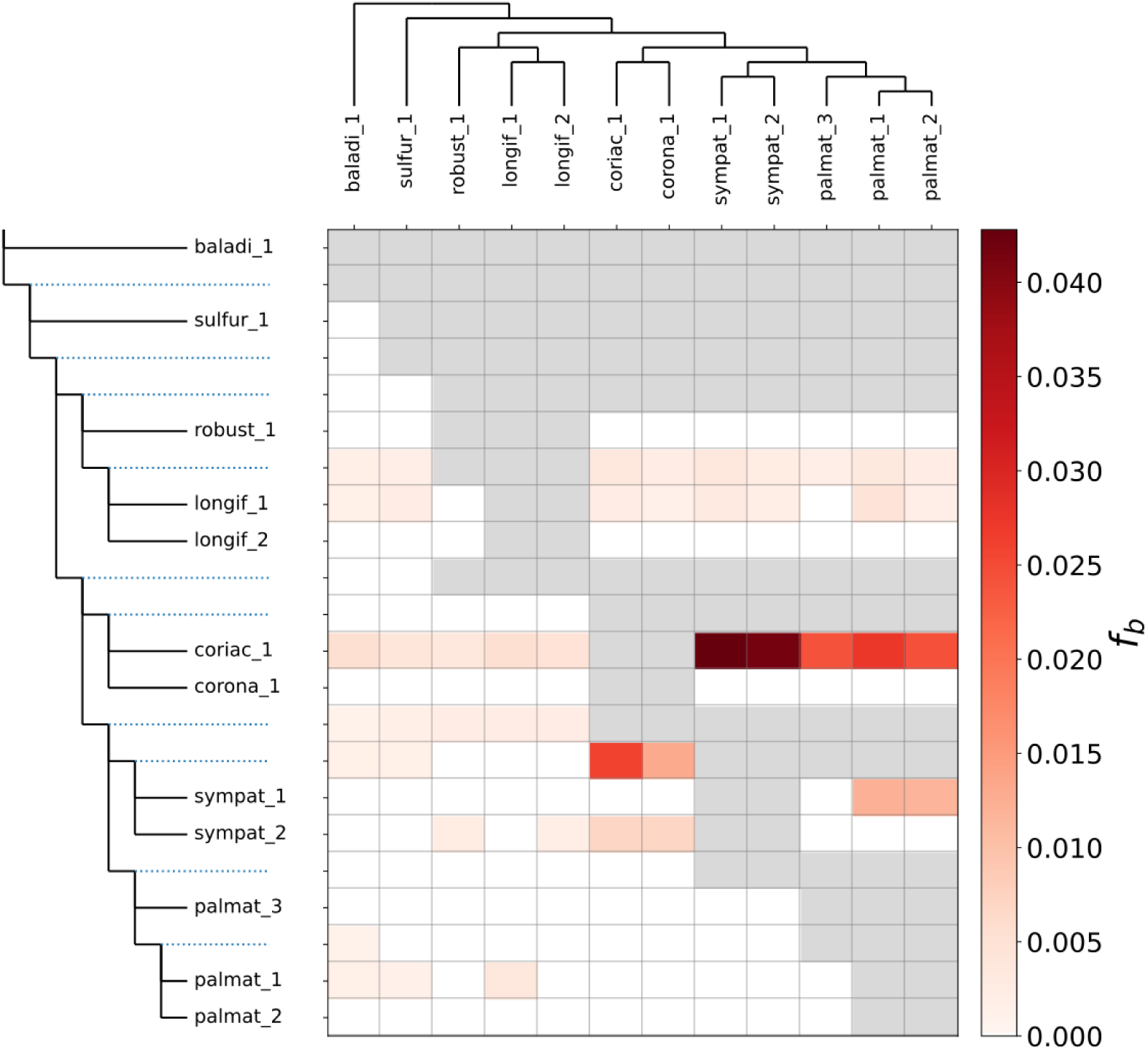
f-branch matrix The f-branch (= fb(P3)) represents admixture proportion between b (y-axis) and P3 (x-axis). The lineage b can take an internal branch which is represented by blue dotted line.

### Population structure analysis

STRUCTURE analysis for *O. palmatinervia*, *O. sympatrica*, and *O. coriacea* showed that delta *K* was the highest at *K* = 3 (supplementary fig.S7). At *K* = 3, *O. palmatinervia*, *O. sympatrica*, and *O. coriacea* had a unique genetic identity illustrated by light blue, dark purple, and orange, respectively (fig. 4). Some individuals of *O. palmatinervia* and *O. sympatrica* from Dzumac and Yate showed limited admixture pattern. At *K* = 2, all of *O. sympatrica* and *O. coriacea* individuals showed admixture of two inferred populations. The population assigned to *O. palmatinervia* (light blue) was dominant in *O. sympatrica* and minor in *O. coriacea*. The degree of admixture was greater in *O. sympatrica* than in *O. coriacea*.

**Fig. 4.**
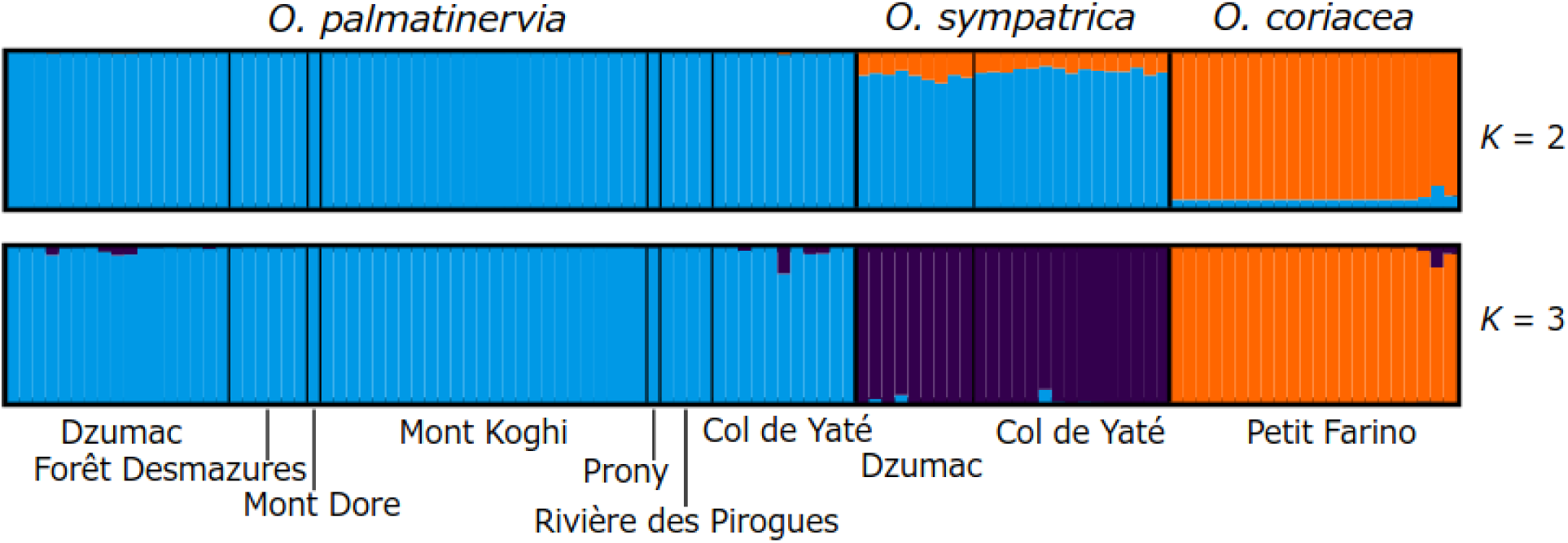
STRUCTURE bar plot based on MIG-seq dataset. Each bar represents one individual, and colored segments represent proportions of genetic components.

**Fig. 5.**
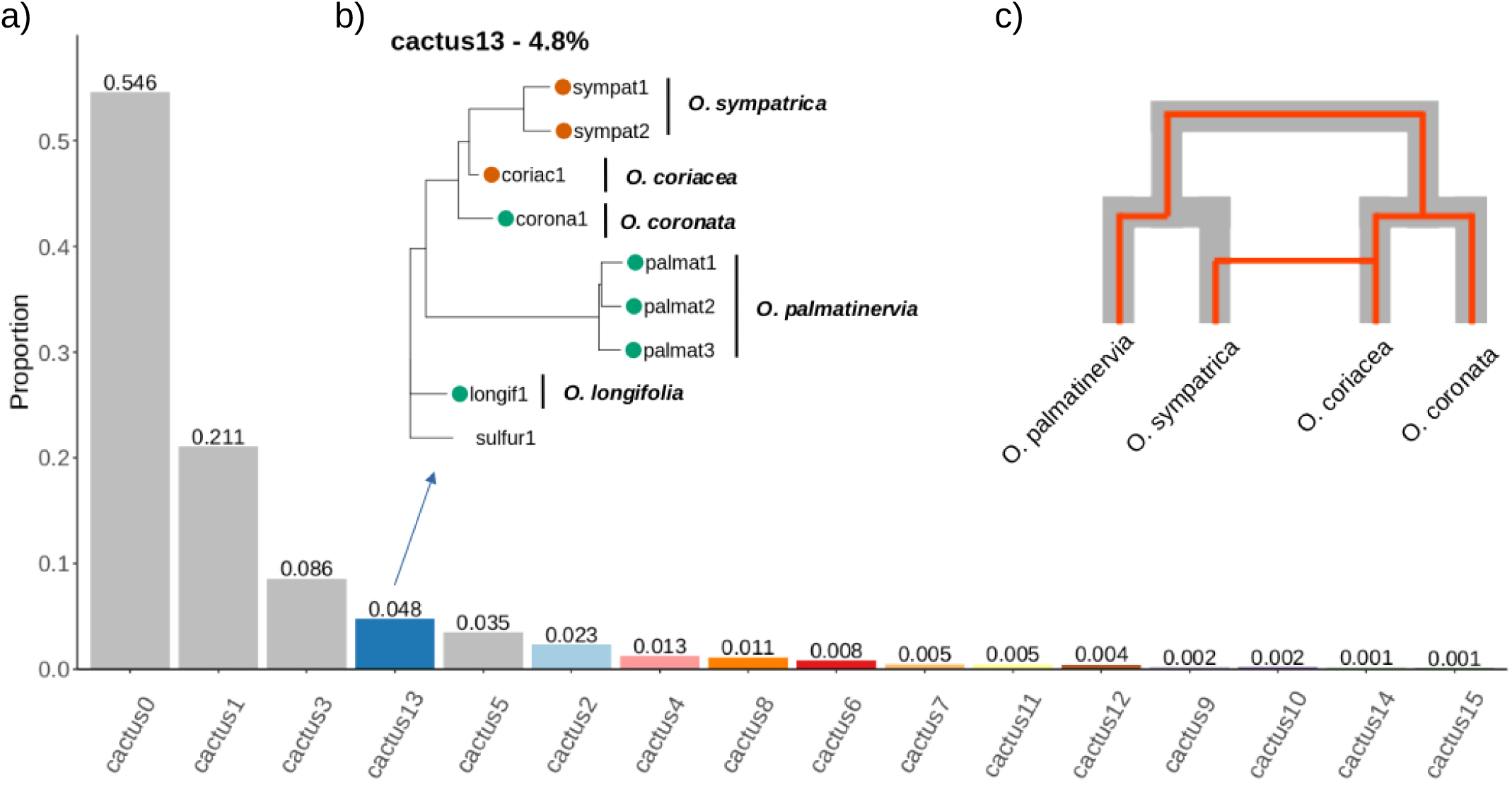
Proportions of each cactus segments, the topology of cactus13 and the expected introgression genealogy. (a) Each bar is colored by the topology of cactus. Grey bar represents the majority topology. (b) The cactus13 is consistent with the expected introgression genealogy. (c) Grey broad line represents genome-wide phylogeny, and red narrow line represents the expected introgression from *O. coriacea* to *O. sympatrica*.

### Introgressed genomic regions from O. coriacea to O. sympatrica

Saguaro and sliding window analysis of fd statistic were used to detect the introgressed genomic regions from *O. coriacea* to the ancestral lineage of *O. sympatrica*. In Saguaro, cactus0, 1, 3, and 5 showed the same topology, and those cactus segments accounted for about 87.8 % of the total segments ( fig.5 and supplementary fig. S9). This major topology is similar to that of estimated phylogenetic trees of IQTREE, SVDQ, and SNAPP based on whole-genome scale SNPs. The cactus13 is highly consistent with an expected introgression genealogy, in which *O. sympatrica* individuals are sister to *O. coriacea* and closely related to *O. coronata* rather than *O. palmatinervia* (fig. 5). The cactus13 segments accounted for 4.8 % of the total segments. Four fd sliding window analyses were performed with different taxon sets. Across all analyses, 153 windows were consistently detected as outliers. In cactus13 segments overlapped with those outlier windows, there are 68 gene records (Supplementary Table S4). This gene list included the MYB gene family and Type II MADS-box gene, as well as the Ovate family protein-coding gene, which encodes proteins that may act as transcription factors.

## Discussion

### Phylogenetic placement of O. sympatrica and its reproductive isolation from the sister species, O. palmatinervia

We inferred phylogenetic relationships within the *robusta* subclade, including a recently described species, *O. sympatrica*, based on whole genome scale SNPs. Our results consistently indicated that *O. sympatrica* is sister to *O. palmatinervia*, with which it coexists sympatrically (fig.1, supplementary fig. S2 and S3). In the STRUCTURE analysis based on MIG-seq data at K = 3, we found that these two species are clearly distinct. It is noteworthy that some samples of *O. sympatrica* and *O. palmatinervia* from Yate and Dzumac, where intimate coexistence (even winding on each other) can be observed, showed weak admixture pattern. This result may reflect relatively recent, perhaps ongoing, localized geneflow between *O. sympatrica* and *O. palmatinervia*. Both phylogenetic inferences and STRUCTURE analysis showed that these two species can be genetically distinguished. It is thought that the reproductive isolation between these two sympatric species is strong enough to maintain their species boundaries.

Floral isolation, a form of pre-pollination reproductive barrier mediated by floral morphology and pollinator behavior (Grant, 1949), is considered to be the primary reproductive barrier between *O. sympatrica* and *O. palmatinervia*. The most pronounced morphological difference between the two species lies in their flower morphology: *O. sympatrica* having pipe-type flowers and *O. palmatinervia* having funnel-type flowers (fig. 1). Field observations suggest that these flower types differ in the pollination efficiency and the pollen placement on the two honeyeater bird species, *Glycifohia undulata* and *Philemon diemenensis*, which are their respective putative pollinators (Gildas *et al*., unpublished data). *Glycifohia undulata*, with its smaller body size, does not come into contact with the elongated stamens and pistils of the funnel-type flower but does contact the shorter stamens and pistils of pipe-type flower well. Conversely, *P. diemenensis*, with its larger body size, was observed to make stamen-pistil contact with both flower types but in a unconventional way for the pipe-type flower. Also, the position of pollen attachment is different between two flower types: pollen from the funnel-type flower was found on the back of their head, while pollen from the pipe-type flower was attached to their beak. In addition, visits by *P. diemenensis* sometimes led to corolla damage in the pipe-type flower, which has a narrower opening than the funnel-type flower. These suggest that the pipe-type flower is specialized for *G. undulata* and the funnel-type for *P. diemenensis*. Although further studies on actual pollen movement by those pollinators and the strength of other reproductive isolation barriers are needed, these mechanical isolations linked to flower shape differentiation likely serves as a major reproductive isolation barrier between *O. sympatrica* and *O. palmatinervia*.

Floral isolation may have played an important role in the early stages of *O. sympatrica*’s speciation process. Our divergence time estimation indicated that *O. sympatrica* and *O. palmatinervia* diverged approximately 0.5 Mya (95% HPD: 0.1-1.0 Mya) (fig.1). The two putative pollinators, *G. undulata* and *P. diemenensis*, are the only New Caledonian representatives of their genera and are endemic to the region (Andersen et al., 2014; Marki et al., 2017). Recent molecular phylogenetic research inferred that *G. undulata* diverged from its Australian and Vanuatuan relatives approximately 5 Mya, while *P. diemenensis* diverged from its relatives in Australia, New Guinea, and Indonesia around 3.5 Mya (Marki et al., 2017; Kumar et al., 2022). Given that the estimated divergence time between *O. sympatrica* and *O. palmatinervia* is considerably more recent than those pollinator divergence times, it is plausible that pollinator environments including *Glycifohia* and *Philemon* species were established in New Caledonia before the speciation of *O. sympatrica*. Adaptation to those pollinators may have driven flower morphological divergence and contributed to the reproductive isolation between *O. sympatrica* and *O. palmatinervia*.

### Ancestral introgression and flower shape evolution in O. sympatrica

Our hybridization analyses indicated that introgression from *O. coriacea* into the ancestral lineage of *O. sympatrica* had occurred. Phylogenetic networks by SnaQ revealed that *O. sympatrica* individuals inherited 11.6 % of the genome from *O. coriacea* (fig.2). Additionally, f-branch statistics detected gene flow between *O. coriacea* and the ancestral lineage of *O. sympatrica* (fig.3). STRUCTURE analysis based on MIG-seq further suggested a bidirectional but uneven gene flow between *O. sympatrica* and *O. coriacea* (fig.4). This ancestral introgression scenario was also supported by genome scan analyses. The Saguaro analysis identified cactus13, which is consistent with the introgression genealogy and the proportion of cactus13-assigned regions was greater than that of other minor cactus-assigned regions (fig.5). In the analysis, there is no cactus which is compatible with the introgression from *O. sympatrica* to *O. coriacea* (fig.s9).

This ancestral introgression may have driven convergent evolution of flower shape in the *robusta* subclade. *O. coriacea* and *O. sympatrica* are the only species with pipe-type flowers in the subclade. Our phylogenetic analysis suggested that flower shape shifts have occurred repeatedly within the subclade (fig.1). Convergent evolution can be driven by single origin mutations, which are shared among distantly related lineages through introgression (Stern, 2013). This pattern of convergent evolution has been showed in the wing color patterns of *Heliconius* butterflies and the red flowers of *Mimulus* monkeyflowers (The Heliconius Genome Consortium, 2012; Stankowski & Streisfeld, 2015). Given the both *O. coriacea* and *O. sympatrica* possess pipe-type flowers, it is conceivable that the ancestral introgression detected in our study may have contributed to the convergent evolution of flower shape in the *robusta* subclade.

We identified introgressed genomic regions shared between *O. sympatrica* individuals from Yate and Dzumac by Saguaro and fd sliding window analysis and then complied a list of genes on them. Transcription factors are known to play crucial roles in plant morphogenesis, including the development of floral organs. Our gene list includes members of the Myb, MADS-box, and OVATE gene families, all of which are potential transcription factors. The Myb transcription factors are thought to contribute significantly to the diversification of pollinator-associated floral traits (Yuan et al., 2013). In *Nicotiana*, overexpression of Myb gene has been shown to result in changes of corolla shape, stamen length and pistil length (Rahim et al., 2019). The MADS-box gene family is well-established for its fundamental roles in floral organ development. In *Brassica rapa*, alternations in the expression of MADS-box genes in the ABCDE model are linked to variations in flower morphology such as pistil length and width (Zhang et al., 2023). Our gene list includes a gene highly similar to the Type II MADS-box transcription factor, which most MADS-box genes in the ABCDE model belongs to. Also, a gene showing high similarity to the ENHANCER OF AG-4, which is involved with the pathway of AGAMOUS, a well-known MADS-box gene related to the ABCDE model, is included. OVATE genes are plant-specific transcription factors that regulate various aspects of growth and development (Wang et al., 2016). In tomatoes, changes in the expression of SlOFP20, a member of the OVATE gene family, have been shown to affect the length of petals, sepals, pistils, and stamens (Zhou et al., 2019). Although the functions of these gene families are diverse and the precise role of each listed gene in *Oxera* remains unclear, it is noteworthy that these genes appear to have introgressed from *O. coriacea* to *O. sympatrica*. Further investigation of selective pressures on these introgressed genomic regions, as well as functional studies of the listed genes, are crucial to elucidating the relationship between the ancestral introgression and the evolution of flower shape in *O. sympatrica*.

Floral traits are classic examples of “magic traits”, which contribute to both environmental adaptation and reproductive isolation (Servedio et al., 2011). When the genetic variants associated with magic traits are introgressed from other species, they can facilitate not only local adaptation but also the evolution of reproductive isolation from closely related species, perhaps even without geographical isolation between them (Jiggins et al., 2008; Salazar et al., 2010; Rosser et al., 2024). The wing color patterns of *Heliconius* butterflies and the flower coloration of *Mimulus* monkeyflowers are well-known instances of such introgression of magic traits. In *Heliconius* butterflies, wing colour patterns are involved in both Müllerian mimicry between toxic species and mate choice as visual markers (Jiggins et al., 2001). In *Mimulus* monkeyflowers, the red flowers, which evolved repeatedly in the genus, are adapted to hummingbird-pollination and play a role in reproductive isolation from yellow-flowered lineages that are pollinated by hawkmoths (Streisfeld & Kohn, 2007; Sobel & Streisfeld, 2015). In *O. sympatrica*, floral isolation may represent a major isolation barrier from its sympatric sister species, *O. palmatinervia*. Hence, if the pipe-type flowers of *O. sympatrica* evolved through the introgression from *O. coriacea*, this ancestral introgression could have played a crucial role in the speciation process of *O. sympatrica*.

In the syngameon hypothesis of adaptive radiation, it is thought that occasional or localized hybridization maintain and prolong the adaptive radiation momentum through the rapid evolution of lineages with new combinations of adaptive traits (Seehausen, 2004). Previous study has shown that in *Oxera*, a high speciation rate has been maintained since the early stage of its diversification (Barrabé et al., 2019). In relatively recent speciations within the subclades of *Oxera*, habitat shift and geographical isolation associated with climate changes during Pleistocene are thought to have been important because their current distributional overlap is quite rare. It is noteworthy that climate changes can cause not only population fragmentation but also secondary contact between diverged populations, providing opportunities for hybridization (Kinoshita et al., 2019; Folk et al., 2023). *O. palmatinervia*, *O. sympatrica* and *O. coriacea* grow in forests on ultramafic soil at approximately the same altitude. However, *O. palmatinervia* and *O. sympatrica* are distributed mainly in the southern part of the island, while *O. coriacea* is distributed mainly in the central part, currently (Gildas *et al*., unpublished data). Distributional changes associated with climate oscillation in Pleistocene may have increased the opportunity for hybridization between ancestral populations of *O. coriacea* and *O. sympatrica*.

The flora of New Caledonia is characterized by a high degree of local microendemism within the island. For their speciation, as with *Oxera*, geographical isolation and local adaptation are thought to have been important (Grandcolas et al., 2008; Paun et al., 2016; Barrabé et al., 2019). In particular, the climate oscillation during Pleistocene and the mosaic of diverse habitats associated with orography, soil, and climate have been highlighted. Those climate changes and the mosaic habitats in New Caledonia may have facilitated the establishment of syngameon and occasional hybridization may have catalyzed the diversification of its flora.

## Conclusions

In this study, we used whole-genome sequencing and MIG-seq to reveal the history of divergence and hybridization within the *robusta* subclade of *Oxera*. Our phylogenetic inferences affirm that *O. sympatrica*, a recently described new species, is sister to *O. palmatinervia* and that convergent evolution of flower shape have occurred in the subclade. Population structure analysis, allele sharing test, phylogenetic network analysis, and genomic local phylogenetic relationships have suggested the ancestral introgression from *O. coriacea* to *O. sympatrica*. Both of these two species have the pipe-type flower and it can be hypothesized that this ancestral introgression resulted in the convergent evolution of flower shape. In the introgressed genomic regions, we found some genes which can be associated with the flower shape development. To our knowledge, there is no study which showed the possibility that introgression may have been involved in the evolution of pollination-related floral traits in New Caledonian plant taxa. Our study has revealed the history of divergence and hybridization of the *robusta* subclade and provided a putative new case of magic trait introgression. Further ecological and genetic studies on *O. sympatrica* and its relatives will provide deep insights into the role of hybridization in the island adaptive radiation and the development of New Caledonia’s rich flora.

## Data availability

The raw sequencing data and the assembled genome data will be deposited in the DDBJ (BioProject ID: PRJDBXXXX) upon publication. All scripts and commands used in this study will be available at https://datadryad.org/dataset/XXXX upon publication.

## Acknowledgments

This work was funded by JSPS Grants-in-Aid for Scientific Research (JP16H06279 (PAGS), JP18H02496, JP21H05314, JP21KK0104, JP22H02699).

