## Supplementary Fig S1-S9 and Table S1-S3 for "Hybridization during the adaptive radiation of *Oxera* (Lamiaceae) in New Caledonia: Is flower shape shift driven by introgression?"

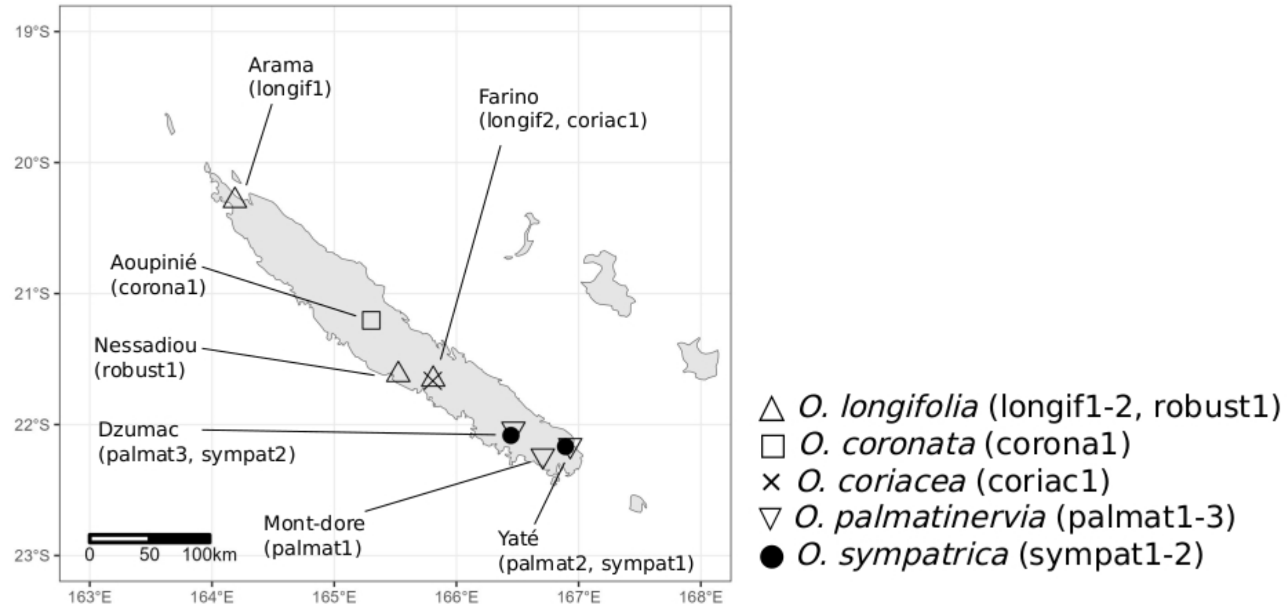


Fig.S1 Sampling sites of the *robusta* subclade individuals for WGS analysis

(a) no-misssing

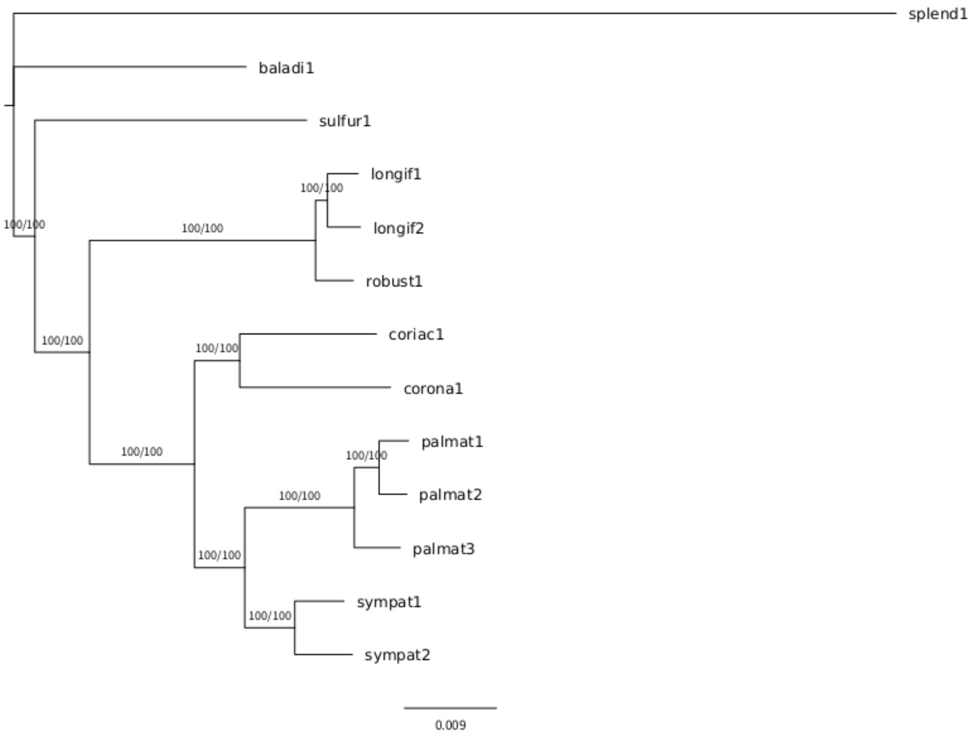


(b) missing < 20%

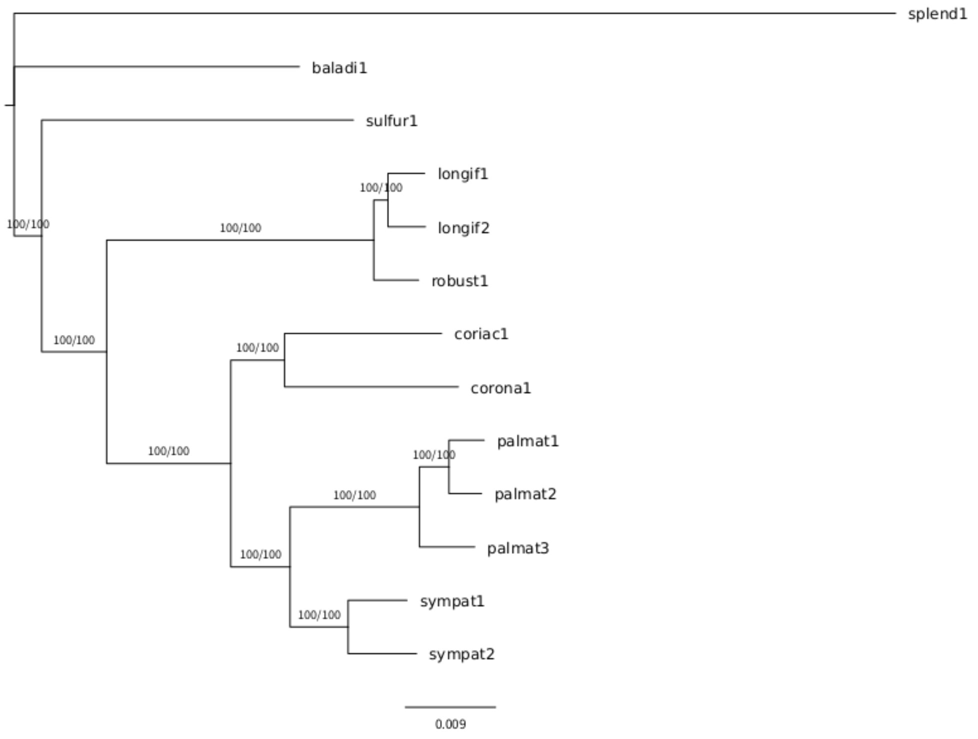


Fig.S2 SNP concatenated ML trees estimated by IQ-tree.
Support values of UFB and SH-aLRT are shown on a branch. (a) no-missing SNPs. (b) missing < 20% SNPs.

(a) 1kb apart

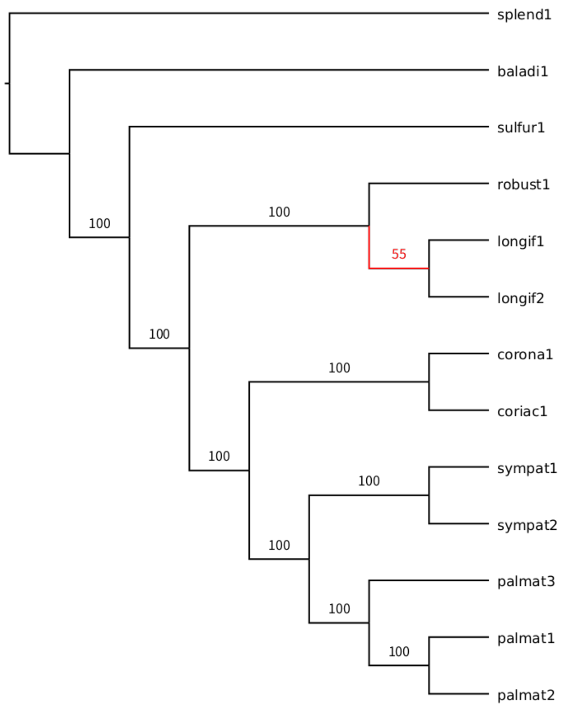


(b) 5kb apart

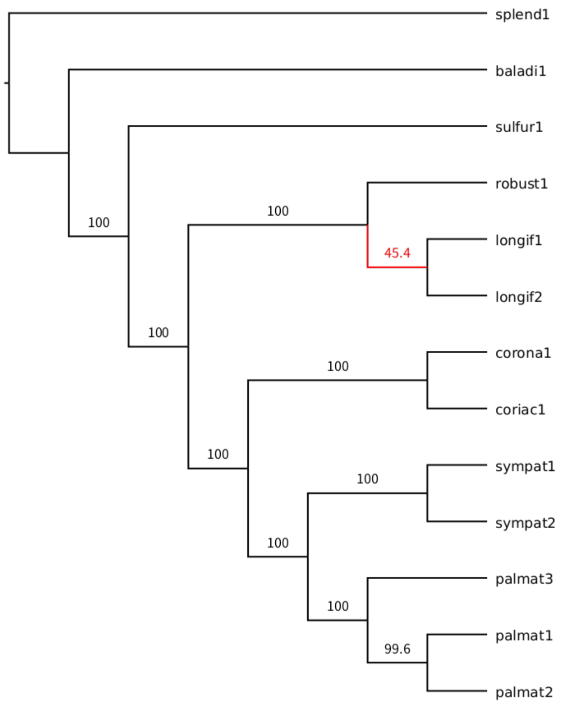


(c) 10kb apart

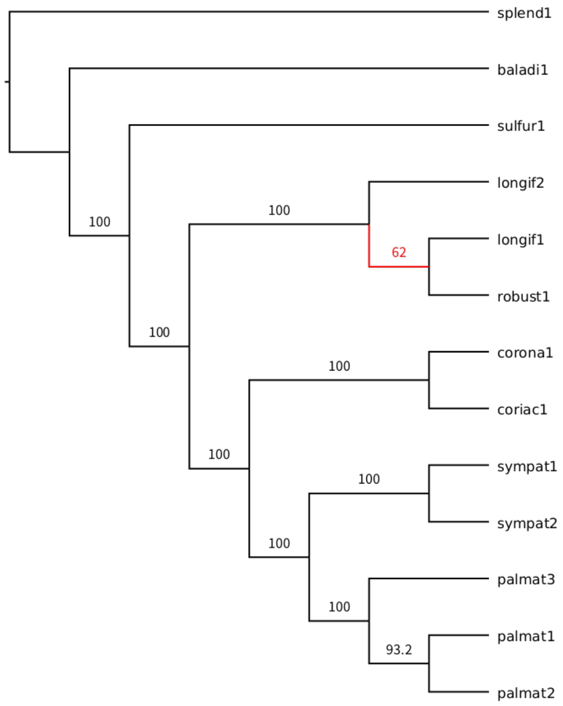


Fig.S3 Multispecies coalescent trees by SVDQ
Support value of bootstrap is shown on a branch. Low support branch is shown in red. (a) 1kb-apart SNP. (b) 5kb-apart SNP. (c) 10kb-apart SNP.

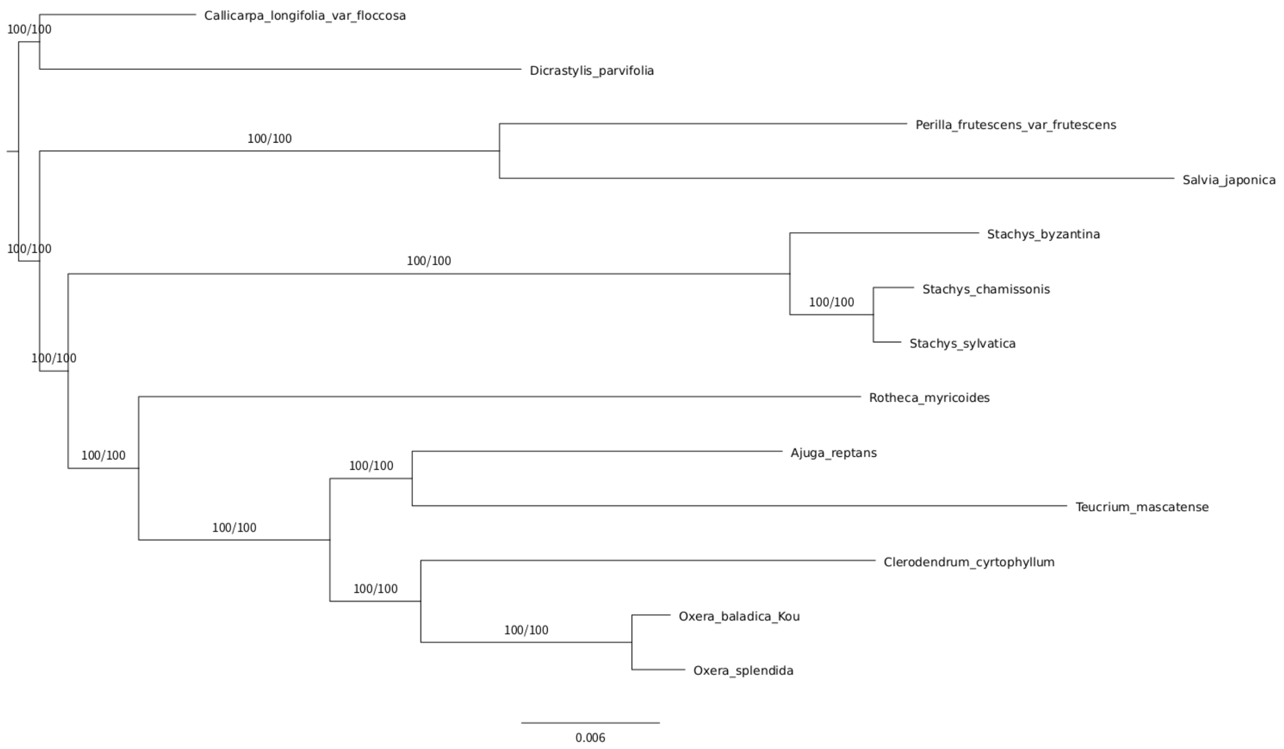


Fig.S4 ML tree of Lamiaceae chloroplast dataset
Support values of UFB and SH-aLRT are shown on a branch.

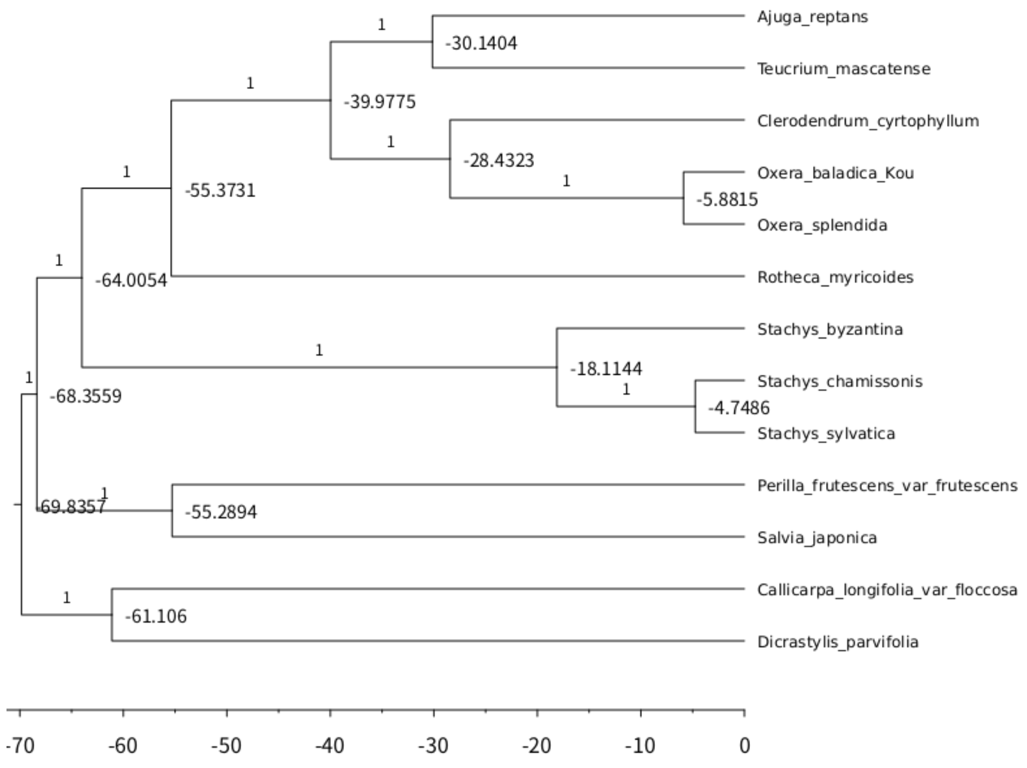


Fig.S5 BEAST MCC tree of Lamiaceae chloroplast genome dataset
Posterior probability is shown on a branch, and mean hight (millions of year) is shown in a node label.

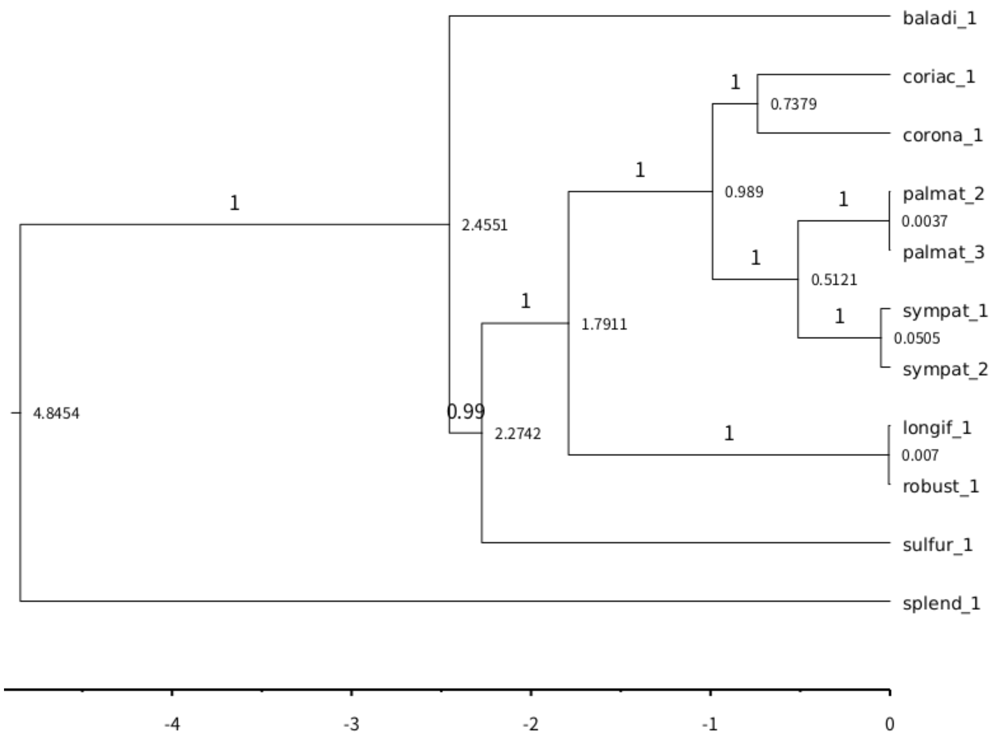


Fig.S6 Time calibrated SNAPP tree based on Genome-wide SNPs

Posterior probability is shown on a branch. Mean hight is shown in a node label.

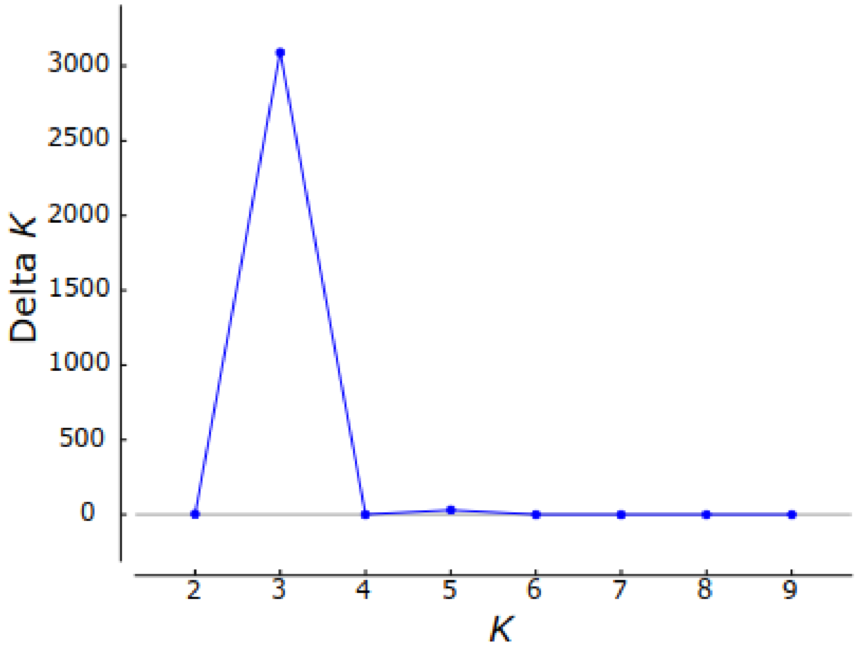


Fig.S7 delta K in STRUCTURE

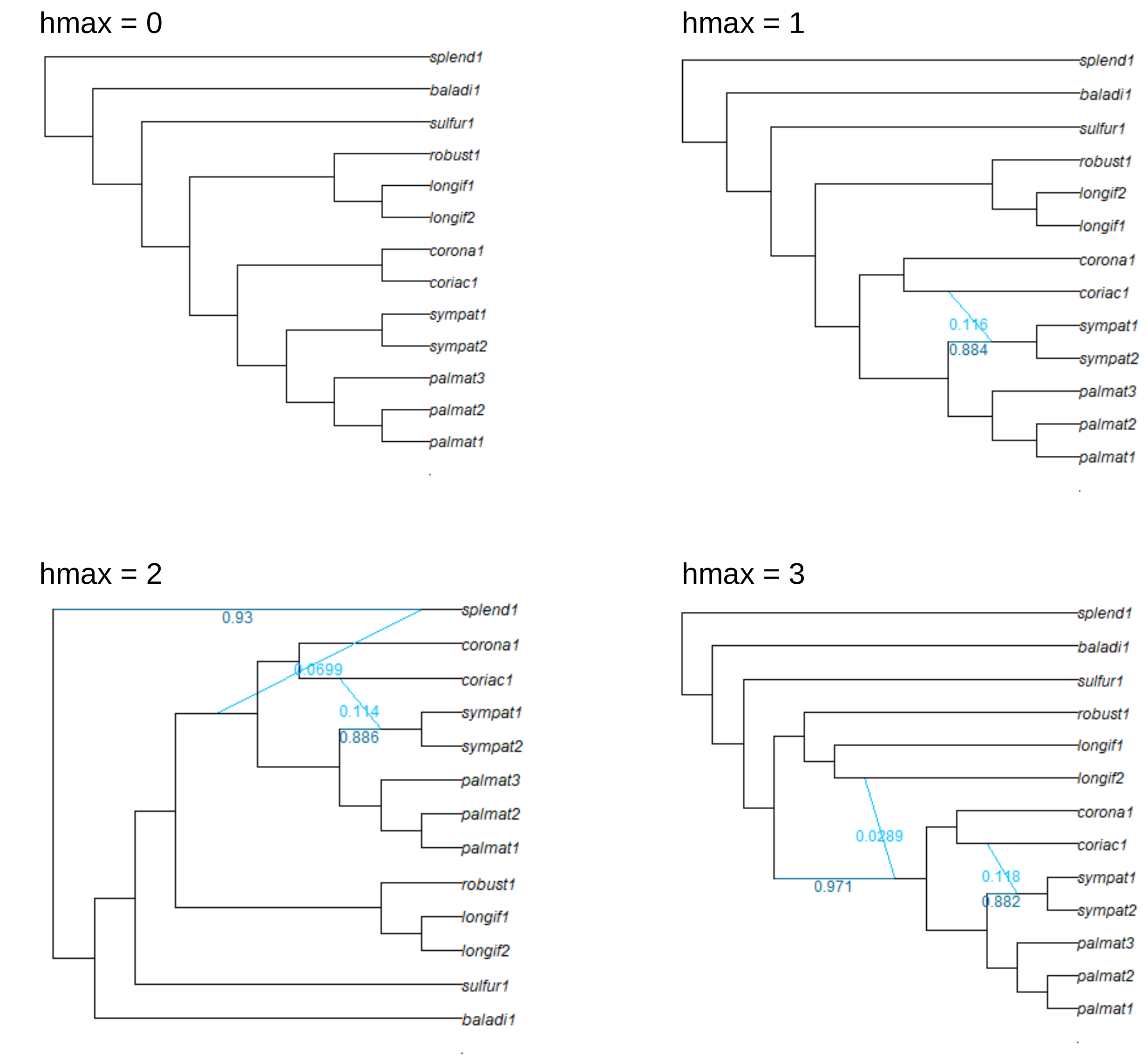


Fig.S8 SnaQ networks based on 2kb-apart SNPs

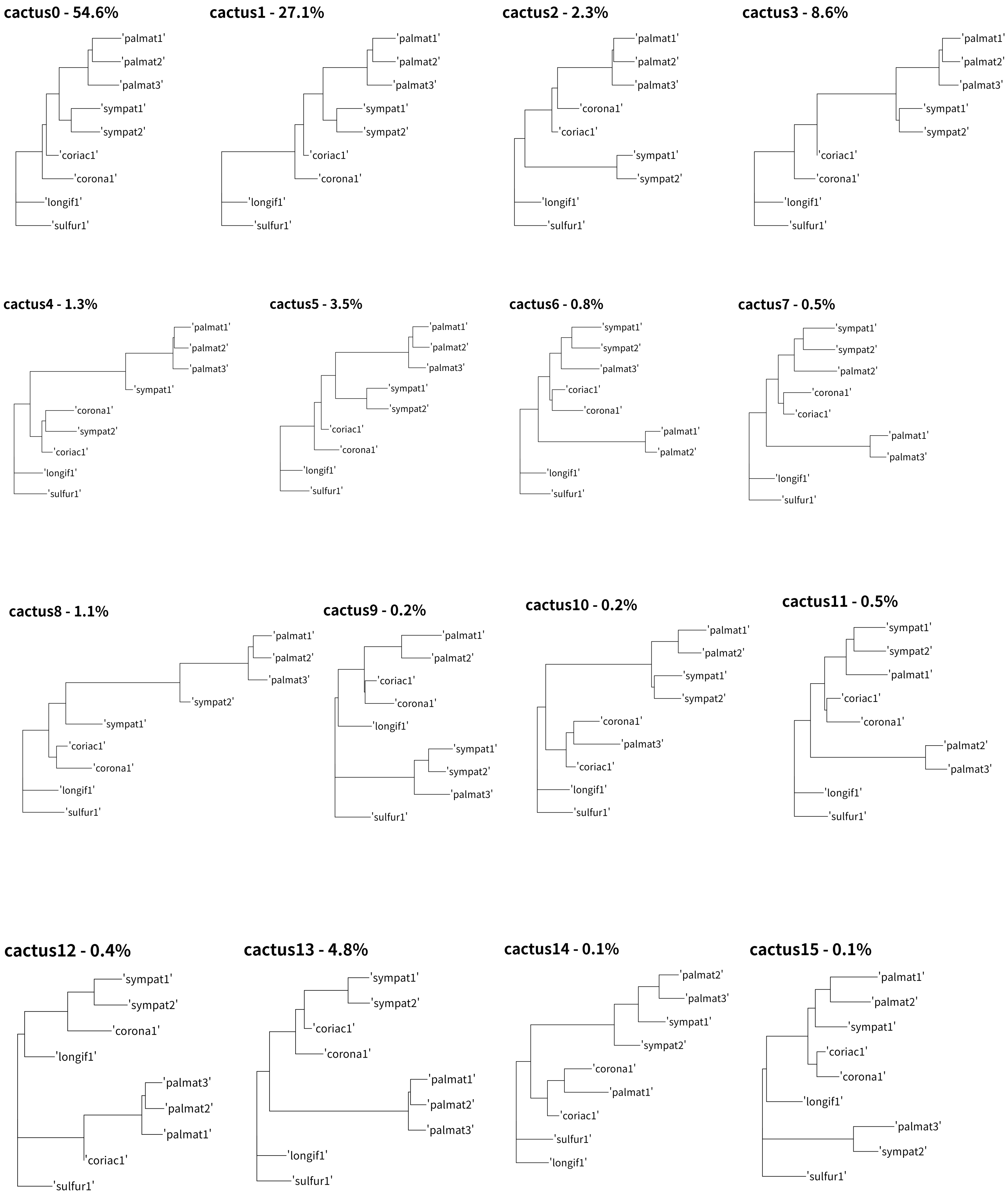


Fig.S9 nj tree of Saguaro cacti

Supplementary Table S1. Sample list of WGS

| sequence name | Species | Location | Sample_ID | DNA_extraction | Library_prep | Sequencing_platform |
|---|---|---|---|---|---|---|
| baladi1 | O. baladica | Koumac, New Caledonia | YI-1351 | CTAB | NEBnext Ultra II | HiseqX |
| coriac1 | O. coriacea | La Foa, New Caledonia | YI-1259 | CTAB | NEBnext Ultra II | HiseqX |
| corona1 | O. coronata | Koumac, New Caledonia | YI-75-C1 | Dneasy Plant Mini Kit | illumina DNA prep | HiseqX |
| longif1 | O. longifolia | Poum, New Caledonia | YI-1088 | CTAB | NEBnext Ultra II | HiseqX |
| longif2 | O. longifolia | La Foa, New Caledonia | YI-1284 | Dneasy Plant Mini Kit | illumina DNA prep | Novaseq 6000 |
| robust1 | O. longifolia | Bourail, New Caledonia | YI-627 | Dneasy Plant Mini Kit | illumina DNA prep | Novaseq 6000 |
| palmat1 | O. palmatinervia | Mont-Dore, New Caledonia | YI-1093 | CTAB | NEBnext Ultra II | HiseqX |
| palmat2 | O. palmatinervia | Yaté, New Caledonia | YI-1317 | Dneasy Plant Mini Kit | illumina DNA prep | Novaseq 6000 |
| palmat3 | O. palmatinervia | Dzumac, New Caledonia | YI-1314 | Dneasy Plant Mini Kit | illumina DNA prep | HiseqX |
| splend1 | O. splendida | Australia | Fa02 (Crayn 1217) | Barrabé et al. (2015) | illumina DNA prep | HiseqX |
| sulfur1 | O. sulfurea | Bourail/Poya, New Caledonia | YI-1347 | CTAB | NEBnext Ultra II | HiseqX |
| sympat1 | O. sympatrica | Yaté, New Caledonia | YI-1315 | CTAB | NEBnext Ultra II | HiseqX |
| sympat2 | O. sympatrica | Dzumac, New Caledonia | YI-1194 | Dneasy Plant Mini Kit | illumina DNA prep | Novaseq 6000 |

Supplementary Table S2. Sample list of Lamiaceae Chloroplast genome dataset

| NCBI Accession number | Species | Subfamily |
|---|---|---|
| MW149076.1 | *Callicarpa longifolia var. floccosa* | Callicarpoideae |
| MT473755.1 | *Dicrastylis parvifolia* | Prostantheroideae |
| KT220689.1 | *Perilla frutescens var. frutescens* | Nepetoideae |
| KY646163.1 | *Salvia japonica* | Nepetoideae |
| KU724141.1 | *Stachys byzantina* | Lamioideae |
| KU724138.1 | *Stachys chamissonis* | Lamioideae |
| KU724140.1 | *Stachys sylvatica* | Lamioideae |
| MW653322.1 | *Rotheca myricoides* | Ajugoideae |
| KF709391.1 | *Ajuga reptans* | Ajugoideae |
| MH325132.1 | *Teucrium mascatense* | Ajugoideae |
| MZ958825.1 | *Clerodendrum cytophyllum* | Ajugoideae |

Supplementary Table S3. Sample list of MIG seq dataset

| Species | Location | number of samples |
|---|---|---|
| *O. palmatinervia* | Dzumac | 17 |
| | Forêt Desmazures | 6 |
| | Mont Dore | 1 |
| | Mont Koghi | 25 |
| | Prony | 1 |
| | Rivière des Pirogues | 4 |
| | Col de Yaté | 11 |
| *O. sympatrica* | Dzumac | 9 |
| | Col de Yaté | 15 |
| *O. coriacea* | Petit Farino | 22 |
