## Supplementary methods for "Hybridization during the adaptive radiation of *Oxera* (Lamiaceae) in New Caledonia: Is flower shape shift driven by introgression?"

### Chloroplast genome assembly and phylogenetic estimation

For chloroplast phylogenetic estimation, chloroplast genomes of *O. splendida* (splend1) and *O. baladica* from Koumac (baladi1) were assembled. *O. splendida* is inferred to be sister to all other *Oxera* species and is distributed outside New Caledonia (Barrabé et al., 2015, 2019). Chloroplast genome assemblies were performed using NOVOplasty 4.2.1 (Dierckxsens et al., 2016) and GetOrganelle pipeline 1.7.6.1 (Camacho et al., 2009; Bankevich et al., 2012; Langmead & Salzberg, 2012; Jin et al., 2020). Assemblies by NOVOplasty were performed from 10,000,000 + 10,000,000 paired reads extracted by SeqKit 0.16.1 (Shen et al., 2016) head command. NOVOplasty was performed with genome range = 130000-160000 and K-mer = 33. The RUBP sequence of *Zea mays* (Gene bank: V00171.1) is used as the seed sequence. Assemblies by GetOrganelle were performed with -R 15, -k 21, 45, 65, 85, 105, and --reduce-reads-for-coverage 800. The chloroplast genome sequence of *O. baladica* was successfully assembled by NOVOplasty and GetOrganelle and a complete sequence match was obtained with those two assemblers. The chloroplast genome sequence of *O. splendida* was successfully assembled by GetOrganelle but could not be assembled by NOVOplasty.

Short reads were mapped to the assembled chloroplast genome sequences and polishing was performed by Pilon 1.24 (Walker et al., 2014). To improve mapping to the end parts of the assembled sequences, the sequences were extended by linking 500 bp of both ends to the other end parts. Sequence extension was performed by SeqKit. Mapping of short reads was performed by BWA 0.7.17-r1188 mem (Li, 2013). Mapped reads were filtered with SAMtools 1.15.1 (Danecek et al., 2021) view -f 2, reads with insert size less than 350 bp were removed with the AWK language, and duplicate reads were removed with GATK 4.1.9 (McKenna et al., 2010) MarkDuplicates with --OPTICAL_DUPLICATES_PIXCEL_DISTANCE 2,500, --REMOVE_DUPLICATES true. The regions corresponding to the original sequence were polished by Pilon based on reads of base quality 20 or higher with --fix snps,indels,gaps, --minqual 20. Base uncertainties were output in IUPAC codes with --iupac. The chloroplast genome sequence of *O. baladica* showed no change after the first polish. The chloroplast genome sequence of *O. splendida* was unchanged after the second polish. Those polished sequences were used for downstream analyses.

The crown age of the genus *Oxera* was estimated based on chloroplast genome and fossil record constraints: the crown ages of the subfamily Nepetoideae and *the Stachys* s.l. clade (Stachydeae excluding *Melittis*). The crown age of the family Lamiaceae has been examined in detail by Rose et al., (2022), and their result was added to the calibration points in our analysis (see below). To include these calibration points in our analysis, the complete chloroplast genome sequences of 11

Lamiaceae species were collected from NCBI, based on the phylogenetic trees of previous studies (supplementary table S2) (Roy & Lindqvist, 2015; Zhao et al., 2021; Rose et al., 2022). The LSC, IR, and SSC regions of each sequence were detected using the organelle genome drawing tool OGDRAW (Greiner et al., 2019) and partitioned using SeqKit. Sequences of each region were aligned by MAFFT 7.508  (Katoh & Standley, 2013) in FFT-NS-i mode with -max-iterate 1000. Then, missing sites (n) were replaced by gaps (-) with the SeqKit replace command. Then, alignment trimming was performed with the trimAl 1.4 (Capella-Gutiérrez et al., 2009) gappyout method. The sequences used for the phylogenetic analysis were 78,510 bp in the LSC region, 25,343 bp in the IR region, and 16,856 bp in the SSC region (total sequence length 120,709 bp).

First, maximum likelihood phylogenetic estimation was performed using IQ-TREE 1.6.12 (Nguyen et al., 2015). The LSC, IR, and SSC alignment data sets were concatenated by catfasta2phyml 1.1.0 (https://github.com/nylander/catfasta2phyml). The branch-linked partition scheme was used (-spp) and substitution models for each region were selected based on BIC using ModelFinder (Kalyaanamoorthy et al., 2017) implemented in IQ-TREE. TVM+F+R2 was selected for the LSC region, K3Pu+F+I for the IR region, and TVM+F+G4 for the SSC region. The UFB and SH-aLRT tests were repeated 1,000 times.

Then, bayesian divergence time estimation was performed using BEAST 2.6.7 (Bouckaert et al., 2019). Base substitution models were set up for each region, while clock model and phylogenetic tree were set up to be shared between the regions. The substitution model for each region was selected by the ModelFinder among the models available in the SSM package 1.1.0 (Walter & Bouckaert, 2018) in BEAST2. In the IR and SSC regions, the substitution models used for IQ-TREE were selected (K3Pu+F+I and TVM+F+G4) and those estimated parameters were used as the initial values for BEAST. In the LSC region, TVM+G4+I was selected. The Optimised Relaxed Clock model (Douglas et al., 2021) was used for the Clock model and the Birth Death model (Gernhard, 2008) for the Tree prior. The following fossil records constraints were used based on the analysis settings of Rose et al. (2022). The subfamily Nepetoideae was constrained as monophyletic and its crown age was constrained by a lognormal distribution (mean in log space = 2.6, standard deviation in log space = 0.5, offset = 47.8). The genus *Stachys* was constrained as monophyletic and its crown age was constrained by a lognormal distribution (mean in log space = 1.5, standard deviation in log space = 0.5, offset = 13.8). In addition to these fossil-based constraints, the crown age of the family Lamiaceae was constrained by a uniform distribution (Lower = 49.95, Upper = 73.2) to include all 95% confidence intervals of estimates in Rose et al. (2022). Besides, based on the phylogenetic trees of previous studies (Rose et al., 2022; Zhao et al., 2021), the most basal divergence of Lamiaceae was constrained as follow: Lineage 1, which includes the subfamilies Callicarpoideae and Prostantheroideae, and Lineage 2, which includes the subfamilies Nepetoideae,

Lamioideae, and Ajugoideae were respectively set to be monophyletic. The divergences of the lineages used for the above constraints were supported by full support values in the phylogenetic estimation by IQ-TREE (supplementary fig. S4).

Three independent runs of Markov chain Monte Carlo (MCMC) with a chain length of 300,000,000 were conducted, sampling every 10,000 generations. MCMC convergence was checked using Tracer 1.7.2 (Rambaut et al., 2018), and the log files and tree files were combined using LogCombiner 2.6.7 with a burn-in of 10%. More than 200 ESSs were identified for all parameters. The Maximum Clade Credibility tree was generated by TreeAnnotator 2.6.7.
